# pyTrance finds co-localizing RNAs in subcellular spatial transcriptomics data

**DOI:** 10.64898/2026.05.07.723470

**Authors:** Leon Strenger, Cledi Alicia Cerda-Jara, Nikos Karaiskos, Nikolaus Rajewsky

## Abstract

Regulation of RNA subcellular localization is crucial for cellular functions in health and disease. For example, local translation of co-localized RNAs is crucial for neural biology. However, it is challenging to identify RNA co-localization events. Here, we present pyTrance, a computational framework that predicts and quantifies subcellular RNA co-localization from spatial transcriptomics data, leveraging latent embeddings learned by a graph neural network. Based on extensive benchmarking, detection of co-localizing RNAs was more accurate and robust compared to existing methods. In mouse brain tissue, pyTrance found several RNA co-localization patterns. Co-localized RNAs were often functionally related and validated by biological knowledge. Interestingly, among novel patterns, pyTrance identified co-localization of GABAergic markers, including *Gad1*, in neuronal projections. Experimental validation led to the discovery of a spatial overlap between *Gad1* mRNA/protein, strongly suggesting local translation. Our results establish pyTrance as a state-of-the-art method to discover biologically important RNA co-localization at subcellular resolution.

## Main

Early examples for the discovery of subcellular RNA localization date back to the 1980s, when it was shown that β-actin is localized to the leading edge of migrating vertebrate fibroblasts (Lawrence & Singer 1986). Since then, numerous additional examples in different organisms and cell types have been discovered, leading to an increasing understanding of the underlying mechanisms, such as intracellular transport and anchoring, as well as the importance of local translation for cell function (Chin & Lécuyer 2017; Das *et al*. 2021; Hollien & Weissman 2006; Mili *et al*. 2008; Moor *et al*. 2017; Wang *et al*. 2016). A prime example of orchestrated RNA localization occurs in neurons, where up to 4000 RNAs have been found to be localized in projections (Perez *et al*. 2021). This allows cells, for example, to produce proteins locally from a single mRNA copy, instead of performing translation in the soma and transporting the resulting proteins to their distant destination (Holt *et al*. 2019; Zappulo *et al*. 2017). Dysregulation of RNA localization or local translation is associated with several neurological diseases; however, the causal relationship is not always fully understood, underlining the importance of further studying RNA localization-related processes (Bauer & Koppers 2025; Engel *et al*. 2020; Fernandopulle *et al*. 2021; Siqueira *et al*. 2025).

Here, we focus on studying the spatial relationship of multiple RNAs (transcribed from the same or different genes). RNA co-localization is usually considered to take place on a nanometer scale, e.g., in the case of mRNA co-translation or in ribonucleoprotein (RNP) granules, which are used by cells, for example, for RNA storage or transport (Cui *et al*. 2024; Shiber *et al*. 2018). Nevertheless, biologically important RNA co-localization can be observed on different scales, from nanometers, like in RNP granules, to several micrometers when investigating RNA in compartmentalized structures, for example, nuclear RNA.

A diverse array of methods is available for studying RNA (co-)localization. These methods usually either perform cell fractionation followed by RNA sequencing (Adekunle & Wang 2020; Benoit Bouvrette *et al*. 2018; Cai *et al*. 2020; Fazal *et al*. 2019; Morf *et al*. 2019) or high-resolution imaging (Alon *et al*. 2021; Rajachandran *et al*. 2025; Scipioni *et al*. 2025; Tsue *et al*. 2024). However, many of these methods are technically complex, often requiring special equipment, which limits their scalability and accessibility across research groups. Additionally, they often have low throughput or are restricted to small gene panels, limiting their use for unbiased discovery of co-localizing RNA groups. More recently, with the advent of high-throughput spatial transcriptomics (ST) methods, RNA localization can be studied unbiasedly at subcellular resolution.

Imaging-based ST (iST) methods are particularly suitable for subcellular RNA localization analysis due to their near single-molecule resolution. Commercially available methods such as Xenium (10x Genomics) (Janesick *et al*. 2023) or CosMx (Bruker Spatial Biology) (He *et al*. 2022) provide easy access to this technology, but the size and complexity of the generated data require advanced computational analysis approaches. However, as the vast majority of these data are aggregated at a single-cell level, few ST analysis tools address subcellular localization. Furthermore, existing approaches mostly come with caveats such as insufficient scalability or reliance on single statistical metrics that do not account for the complexity required to model co-localization of many RNAs across cells (Bierman *et al*. 2024; Kumar *et al*. 2024; Mah *et al*. 2024; Walter *et al*. 2023; Wang & Zhou 2025). Moreover, most of them are designed for analyzing localization patterns of individual RNAs and not co-localization patterns.

To overcome these problems, we present pyTrance, a **Py**thon framework for **tran**script **c**o-localization analysis based on latent **e**mbeddings for iST data. pyTrance trains a graph neural network (GNN) based on each cell’s spatial transcript neighborhood graph in a self-supervised fashion. The computed low-dimensional transcript embeddings are aggregated to gene embeddings, which encode their transcriptional spatial neighborhoods. Assuming that co-localizing RNAs have similar neighborhoods, clustering on the gene embeddings reveals different co-localization groups. To statistically validate co-localization groups, pyTrance introduces an adjusted version of the colocation quotient (CLQ) that accounts for random localization (Leslie & Kronenfeld 2011). Versions of the CLQ have already been applied for analyzing ST data on cellular (Bouchard *et al*. 2025) and subcellular (Mah *et al*. 2024) levels, but our adjustment renders it interpretable and comparable across cells and thus allows ranking cells by the co-localization strength of a given group and identifying outlier populations. The GNN-based approach allows pyTrance to detect co-localization groups of variable sizes, compared to, e.g., distance-based metrics, which have to be recomputed for each combination of genes and usually work only for gene pairs. Furthermore, by sharing gene weights and thereby their co-localization information across cells, the model can detect global patterns more robustly, also for genes with low expression.

We evaluated pyTrance’s ability to detect co-localizing RNAs in simulated and real datasets and compared it to existing methods in an extensive benchmark, in which pyTrance-derived clusters recapitulated the true localization patterns more accurately. In a mouse brain dataset, we predicted Gad1, Gad2, Kcnmb2, and Hapln1, four GABAergic neuron markers, to co-localize. We experimentally validated these predictions using single-molecule fluorescence in situ hybridization (smFISH) in tissue and cultured neurons, where we found the four RNAs to co-localize in neuronal projections. Using additional stainings for the corresponding Gad1 protein showed that its RNAs and proteins co-localize outside of the soma, which suggests that the RNAs are translated locally.

## Results

### The pyTrance workflow

pyTrance identifies co-localizing genes through a three-part workflow designed for subcellularly resolved iST data in count-based format, i.e., each spot corresponds to one detected transcript (**Fig. 1a**). First, pyTrance trains a GNN that takes the transcript-by-gene expression matrix and a spatial transcript neighborhood graph as input for each cell. For the results reported in this study, we constructed the graph based on a radius within which a spot is connected to all other spots. This has the advantage that the radius controls the spatial scale on which co-localizations are detected. Alternatively, a k-nearest neighbors approach is implemented as well. Based on the expression matrix and neighborhood graph, the GNN computes an embedding with reduced dimensionality for each spot, which reflects its transcriptional neighborhood. The model is trained in a self-supervised fashion by differentiating original inputs from permuted copies, forcing it to learn co-localization patterns only found in the original inputs (Methods) (Veličković *et al*. 2018). Here, we use a GNN with only a single graph convolutional layer to ensure that information is passed only within the user-defined radius (Kipf & Welling 2016).

**Fig. 1:**
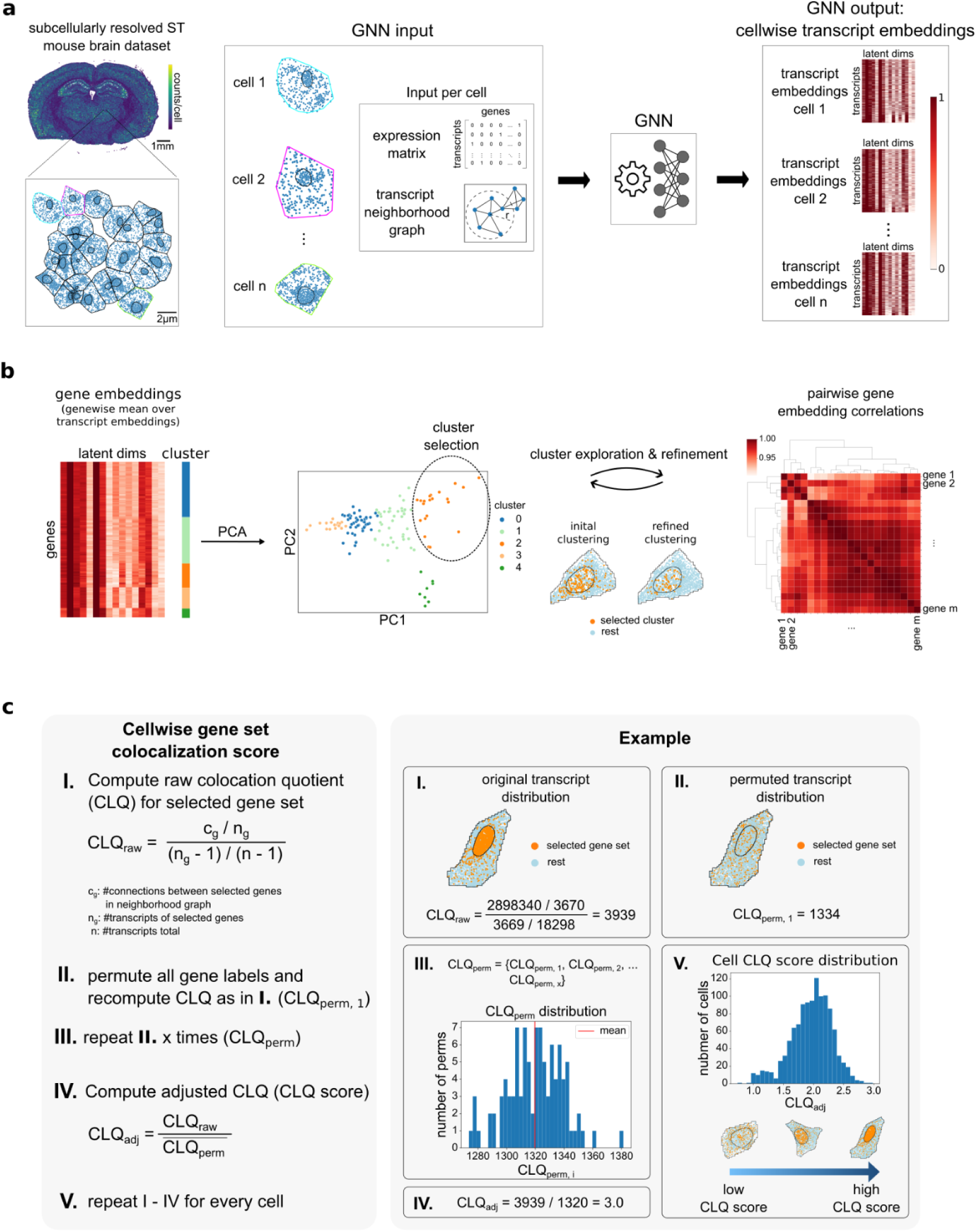
pyTrance Workflow. **a**, Overview of the Graph Neural Network (GNN) processing pipeline, left: Example ST data colored by total counts per cell, single transcripts shown as blue dots in zoom-in; middle-left: GNN input consisting of expression matrix and neighborhood graph per cell; right: computed low-dimensional transcript embeddings per cell. **b**, Transcript embeddings are aggregated for each gene by computing the mean of its transcript embeddings, resulting gene embeddings are clustered by similarity and reduced to two dimensions for visualization using PCA, resulting clusters are manually explored and potentially refined. **c**, A co-localization score for a selected set of genes is computed for each cell, indicating strength of co-localization for the corresponding transcripts.

To find co-localizing RNAs across cells, pyTrance computes the mean over all transcript embeddings corresponding to the same gene, yielding gene embeddings, which are subsequently clustered (**Fig. 1b**). The resulting clusters are treated as predictions of RNA co-localization and can be explored through principal component analysis (PCA), pairwise embedding correlations to identify potential subclusters, as well as visualization of the actual subcellular spatial distribution of the corresponding transcripts. The clustering can be refined iteratively to the desired level of granularity.

As a last step, we validate the co-localization of a selected set of genes statistically using the colocation quotient (CLQ) (**Fig. 1c**) (Leslie & Kronenfeld 2011). Compared to metrics based purely on counting connections or distance measures, the CLQ has the advantage that it takes the number of transcripts of the gene set as well as total transcripts in the cell into account, which makes it more suitable for comparing results of gene sets with different expression levels within the same cell. However, due to the metric’s count-based nature, the resulting values are neither comparable across cells with different expression levels nor interpretable in terms of co-localization strength. Therefore, we initially compute the CLQ based on its original definition (“raw CLQ”) and adjust it for random localization (“adjusted CLQ”), similar to (Bouchard *et al*. 2025)) (Methods). The resulting value, here called “CLQ score”, can be interpreted as quantifying the observed co-localization strength over the expected (random); e.g., a value of “2” indicates that the transcripts from the gene set co-localize twice as often as would be expected if transcripts were randomly distributed within the cell, given the overall spatial distribution of spots. Thus, the CLQ score can be used to quantify the co-localization strength of a gene set across cells, as well as to identify subsets of cells with stronger or weaker co-localization. pyTrance is available as an open-source Python package and incorporates the GNN workflow for unbiased prediction of RNA co-localization as well as quantification through the CLQ score.

### PyTrance accurately identifies co-localizing genes in simulated data

To validate pyTrance, we created a simulated dataset using the Sim-FISH framework (Imbert *et al*. 2022). We simulated the expression and subcellular localization of 450 genes, which corresponds to a medium-sized iST gene panel, following six different localization patterns implemented in Sim-FISH with varying localization intensity in 5000 cells (**Fig. 2a**, **Extended Data Fig. 1a-d**, Methods). Additionally, the genes were split into three groups with different expression levels. We expected to identify the different patterns using the clustered gene embeddings computed with pyTrance, as transcripts following the same pattern localize in the same cell compartments. This compartmental co-localization does not hold for single foci genes, which form focal aggregates independently from one another. Indeed, four of the obtained gene clusters resemble the nuclear, nuclear edge, cell edge, and random patterns, with only foci and cytoplasm genes being mostly mixed in two clusters (**Fig. 2b, c**).

**Fig. 2:**
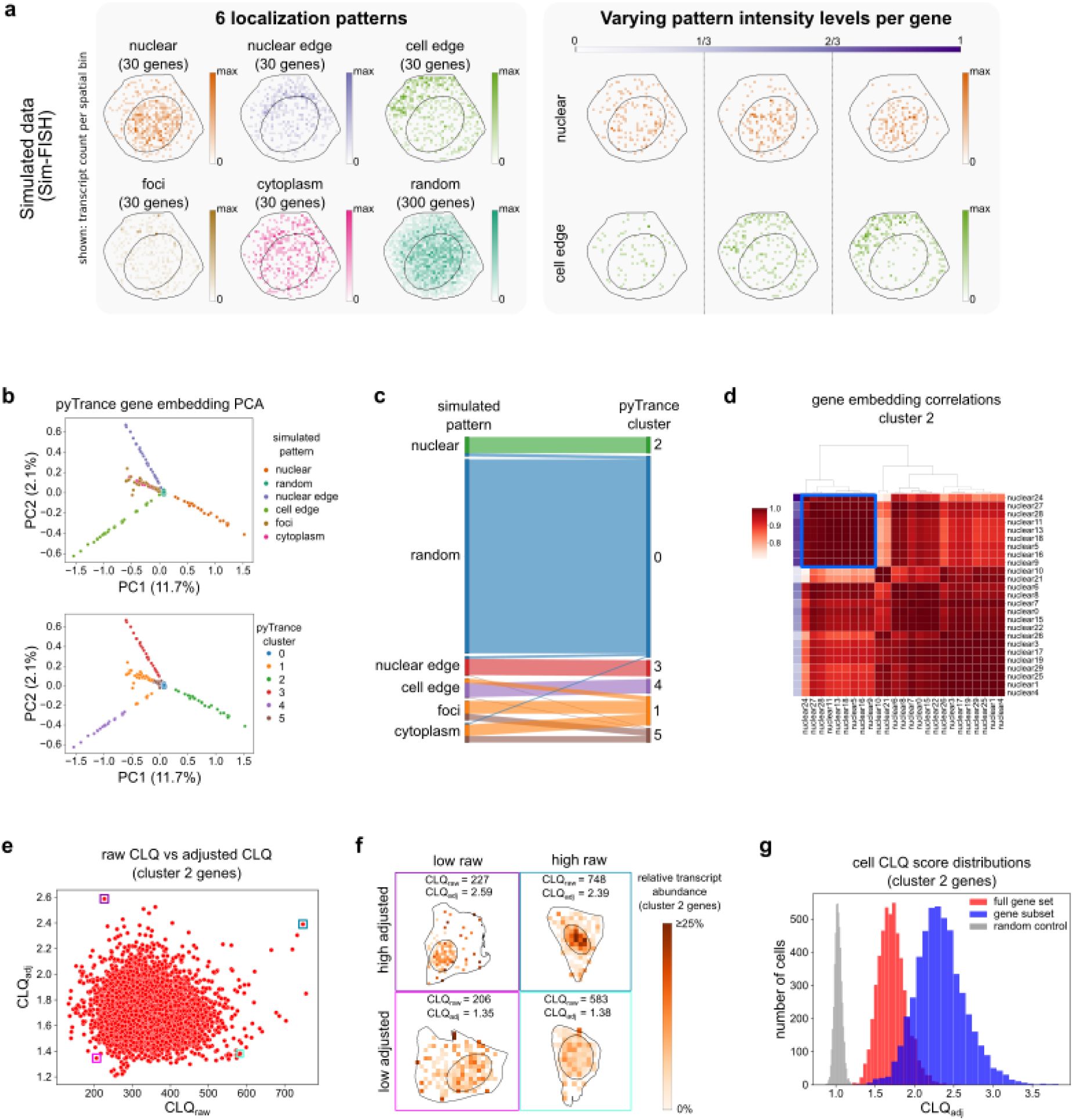
pyTrance detects simulated co-localization patterns in simulated data. **a**, left: The six simulated localization patterns in an example cell. All expressed genes are shown cumulatively. Right: Examples of different localization intensities for two patterns. All genes within each interval are shown cumulatively. In each plot the single transcripts are binned and summed per bin with different maximum values for each plot. **b**, PCA of the gene embeddings computed with pyTrance colored by the true simulated patterns (top) and clusters obtained from the embedding space (bottom). **c**, Comparison of simulated gene patterns and computed gene colocalization clusters. **d**, Pairwise Pearson correlation matrix of gene embeddings of cluster 2. The purple gradient indicates the simulated localization intensity. The blue square indicates a subset selected based on high correlation. **e**, Raw vs adjusted CLQ of genes in cluster 2 for each cell. **f**, Example cells with high/low raw and adjusted CLQs highlighted in **e**. Shown is the transcript count of the selected genes relative to the total counts per spatial bin (relative transcript abundance). **g**, Distribution of adjusted cell CLQs for all cluster 2 genes (red), the subset with high embedding correlations highlighted in **d** (blue), and a set of genes with random localization (grey).

The simulated gene pattern intensities within the obtained clusters showed multiple trends (**Extended Data Fig. 1e**). First, the genes in cluster 5, as well as nuclear, nuclear edge, and cell edge genes assigned to the wrong cluster, all have very low intensities close to a random distribution, making them hard to distinguish. Furthermore, the subclusters we found in the gene embedding correlation heatmaps of clusters 2, 3, and 4 are highly concordant with the genes’ pattern intensities, showing that pyTrance is able to detect different degrees of co-localization strength independent of the genes’ expression levels (**Fig. 2d**, **Extended Data Fig. 1f**). Lastly, most of the foci genes in cluster 1 with high intensity show comparatively low correlation with the rest of the cluster, indicating the GNN was in fact able to detect their localization patterns that are independent for each foci gene.

Next, we compared raw and adjusted CLQ regarding their ability to score cells for co-localization strength of a selected gene set. We computed both metrics on the genes of cluster 2 across all cells and found no correlation between the two metrics (Pearson correlation coefficient: −0.07) (**Fig. 2e**). In cells with low or high values for both metrics in their respective range, only the adjusted CLQ separated cells with different co-localization strengths reliably (**Fig. 2f**, **Extended Data Fig. 2a**). This can be largely explained by the differences in cell size and transcript density, which strongly affect the raw CLQ due to its count-based nature (**Extended Data Fig. 2b**). The negative correlation of the adjusted CLQ and transcript density can be explained by the lower relative nucleus sizes with respect to the whole cell, which leads to a more confined localization for the nuclear genes of cluster 2. The advantage of the CLQ adjustment is further confirmed when comparing both metrics’ value distributions for: (1) the full set of genes of cluster 2, (2) a control group of genes with a random pattern, and (3) a subset of genes of cluster 2 with high intensity (**Fig. 2g**, **Extended Data Fig. 2c**). Especially for the random-patterned genes, our CLQ score distribution shows a clear separation and is centered at 1, underlining the better interpretability. We confirmed that cells with higher scores showed a stronger co-localization for the full gene set. In addition, the gene subset showed a stronger co-localization in the same cells, as suggested by the higher scores and intensities (**Extended Data Fig. 2d**).

Taken together, our analyses show that pyTrance reliably detects and quantifies RNA co-localization for different spatial patterns and localization intensities in simulated data.

### Comparison to other methods

We benchmarked pyTrance against currently published methods for ST subcellular localization analysis, including Bento (Mah *et al*. 2024), InSTAnT (Kumar *et al*. 2024), FISHfactor (Walter *et al*. 2023), Sprawl (Bierman *et al*. 2024), ELLA (Wang & Zhou 2025), and CellSP (Aggarwal & Sinha 2025), where the Bento package consists of three independent methods (RNAflux, RNAforest, and RNAcoloc). First, we compared the methods based on five categories: (1) scalability, which we further divided into runtime and memory consumption; (2) their ability to detect co-localization of RNA transcribed from different genes (i.e., variable numbers of genes); (3) whether the methods are trained in a self- or unsupervised fashion; (4) their ability to leverage co-localization information across cells; and (5) whether they can predict or score pattern labels directly without manual inspection by the user (Table 1). CellSP did not return any results and was therefore not considered further (Methods).

1. To compute runtime and peak memory usage, the methods were run with default parameters on the full simulated dataset (Methods). RNAforest had the best overall scalability, while pyTrance was among the top three methods. We note that most of pyTrance’s runtime is due to training the GNN, which has almost converged to peak performance after ∼5% of the total epochs, corresponding to a runtime of ∼2 hours (**Extended Data Fig. 3a**).
2. RNAcoloc, InSTAnT, and Sprawl are all unable to detect co-localizing groups of more than two RNAs. While Sprawl computes pattern scores for each gene individually and is therefore not designed to detect co-localization out of the box, RNAcoloc and InSTAnT both compute co-localization for pairs of two genes. The embedding-clustering approach of pyTrance allows for variable group sizes, including single-gene clusters for genes with distinct patterns.
3. The only method trained in supervised fashion, hence requiring annotated training data, is RNAforest. It comes already pretrained such that it can be readily applied to new datasets without the need for further annotated data; however, this limits its predictions to patterns it was trained on. pyTrance requires no annotations due to its self-supervised GNN training.
4. Next to pyTrance, only FISHfactor and ELLA share information across cells, which can improve performance, especially in sparse datasets and for lowly expressed genes.
5. For methods designed to find co-localizing genes, including pyTrance, assigning a pattern label to the obtained co-localization groups requires manual inspection by the user. Only RNAforest and Sprawl compute predictions for predefined localization or pattern types. Even though ELLA also predicts localizations for each gene, it does so purely based on distance from the cell center and does not model different types of patterns.

**Table 1:**
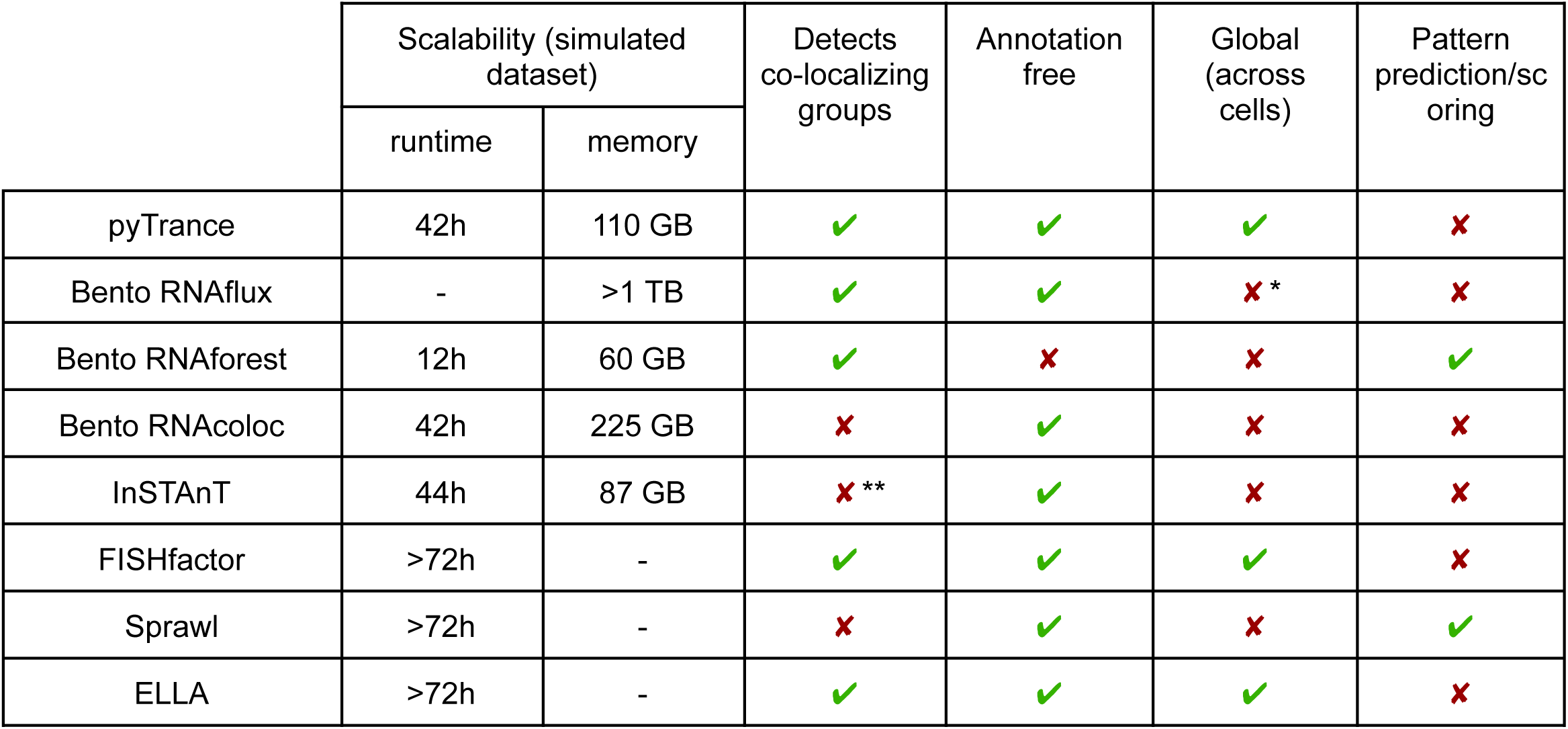
Comparison of published sub-cellular localization analysis methods. ‘Detects co-localizing groups’: groups of variable size. ‘Annotation free’: a method does not require an annotated dataset to be trained. ‘Global’: a method shares information of gene localization across cells to compute its predictions. Considered were peer-reviewed methods designed for ST data. RNAflux, FISHfactor, Sprawl and ELLA could not be run due to excessive memory usage or runtime. *RNAflux applies dimensionality reduction of embeddings across cells, but the embeddings are computed for each cell fully independently. **InSTAnT provides a function to group genes based on pairwise co-localization, however, we were not able to obtain results from that.

Next, we compared the existing methods to pyTrance by their ability to cluster the genes by their simulated localization patterns, as demonstrated above for pyTrance. To do so, we systematically subset the dataset by applying thresholds on the number of cells, minimum pattern intensity levels of genes, and total number of transcripts (Methods). Since RNAcoloc and InSTAnT only compute pairwise co-localizations and FISHfactor, Sprawl, and ELLA did not scale to the size of the dataset, only RNAflux and RNAforest could be included in this benchmark. The performance was measured using two different metrics that quantify the similarity between true gene labels and predicted clusterings: the adjusted Rand index (ARI) and the adjusted mutual information (AMI). For RNAflux, we assigned the genes to clusters based on their highest fluxmap values, as the developers suggest interpreting the fluxmaps as different localization patterns. However, results from this approach were very poor, indicating that fluxmaps cannot be assigned to single compartments across cells (**Extended Data Fig. 3b**). Treating the fluxmaps as embeddings and clustering them, similar to the approach we use for pyTrance, improved results significantly (**Extended Data Fig. 3c**).

Overall, we found that pyTrance performed better on almost every data subset according to the ARI score, while the AMI score of RNAflux was similar on many subsets (**Fig. 3, Extended Data Fig. 3d**). RNAforest appeared to perform better than RNAflux on most data subsets according to the ARI score. However, at closer inspection, we found that in many cases RNAforest misclassified whole groups of genes, e.g., “nuclear edge” genes were predicted to be “nuclear”, which is not necessarily penalized by the evaluation metrics, as they treat the predictions as clusters instead of class labels (**Extended Data Fig. 3e**). This was also the case for the data subsets with only 10% of the transcripts, where RNAforest appeared to perform better than pyTrance (**Extended Data Fig. 3f**).

**Fig. 3:**
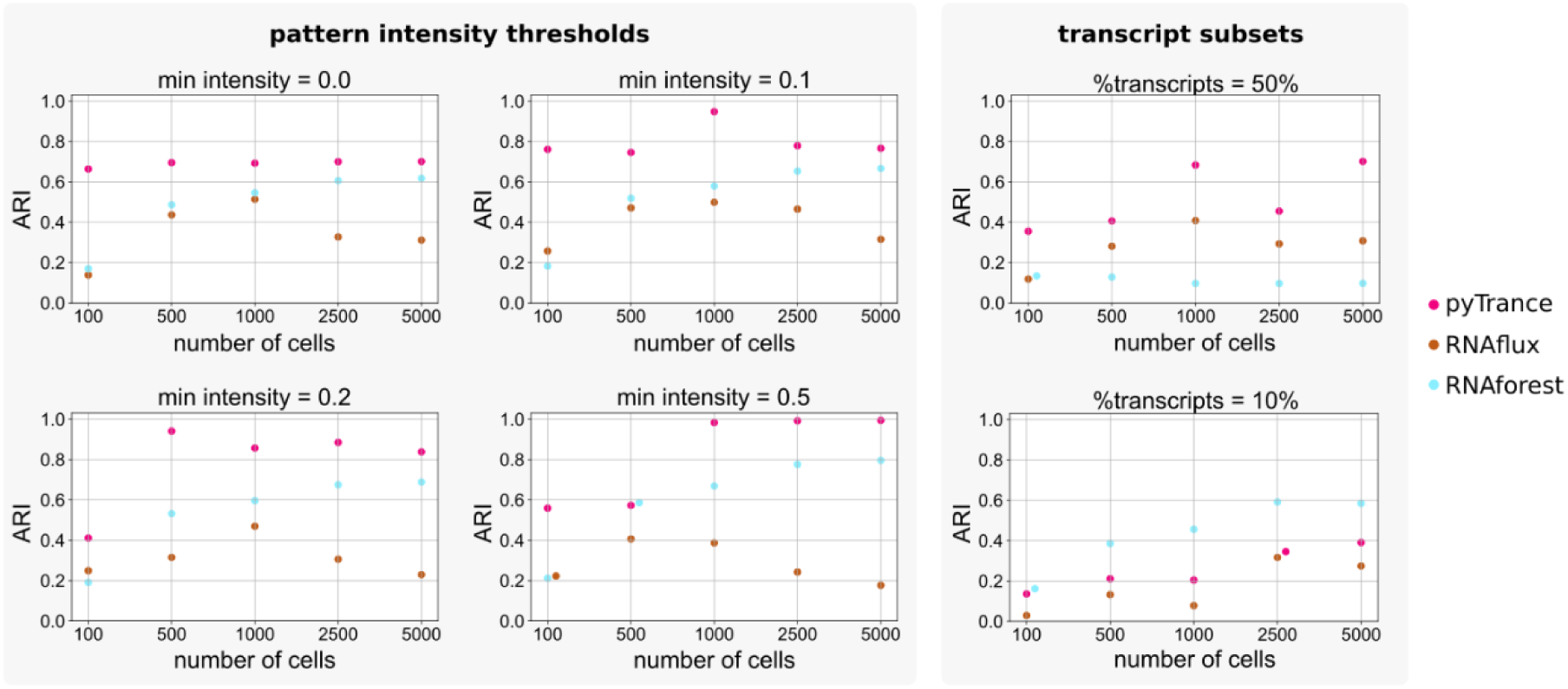
Performance benchmark of pyTrance, RNAflux and RNAforest on detecting simulated co-localizing genes. The full simulated dataset was downsampled in the number of cells (x-axis). Subsets were either filtered for genes with minimum simulated pattern intensities (left) or percentages of the total number of transcripts were sampled (right). ARI was computed how well the computed clusters (pyTrance and RNAflux) or predicted labels (RNAforest) match the simulated patterns.

Moreover, we compared the results of three different clustering algorithms for the pyTrance embeddings and found that k-means clustering typically produced higher scores (**Extended Data Fig. 4a**). However, the Leiden resolution parameter was fixed to avoid subset-dependent fine-tuning, which resulted in varying numbers of clusters across subsets for both pyTrance and RNAflux (**Extended Data Fig. 4b**). This explains the score fluctuations for both methods and shows that the performance could be improved with a more careful clustering parameter selection (**Extended Data Fig. 4c**). We therefore recommend users to explore different clusterings manually when analyzing a new dataset, especially because cluster numbers and boundaries are usually (for “real” data) not as clearly defined as in the given simulated data.

In summary, we found that pyTrance fills a gap in the landscape of subcellular RNA localization analysis methods by providing a scalable and flexible framework that outperforms the comparable existing methods in detecting co-localizing RNAs in simulated data.

### pyTrance co-localization analysis identifies dividing cells in U2OS cell line data

Next, we applied pyTrance to an immortalized human cancer cell line dataset (U2OS cells) generated with the MERFISH technology (**Fig. 4a**) (Mah *et al*. 2024). Cell line data have the advantage that they do not suffer from z-stack noise frequently observed in tissue data, where cells overlap in 3D (Tiesmeyer *et al*. 2026). Furthermore, the transcript density in the used cell line data is even higher than in our simulated dataset, rendering it an ideal scenario to evaluate pyTrance’s performance on real data.

**Fig. 4:**
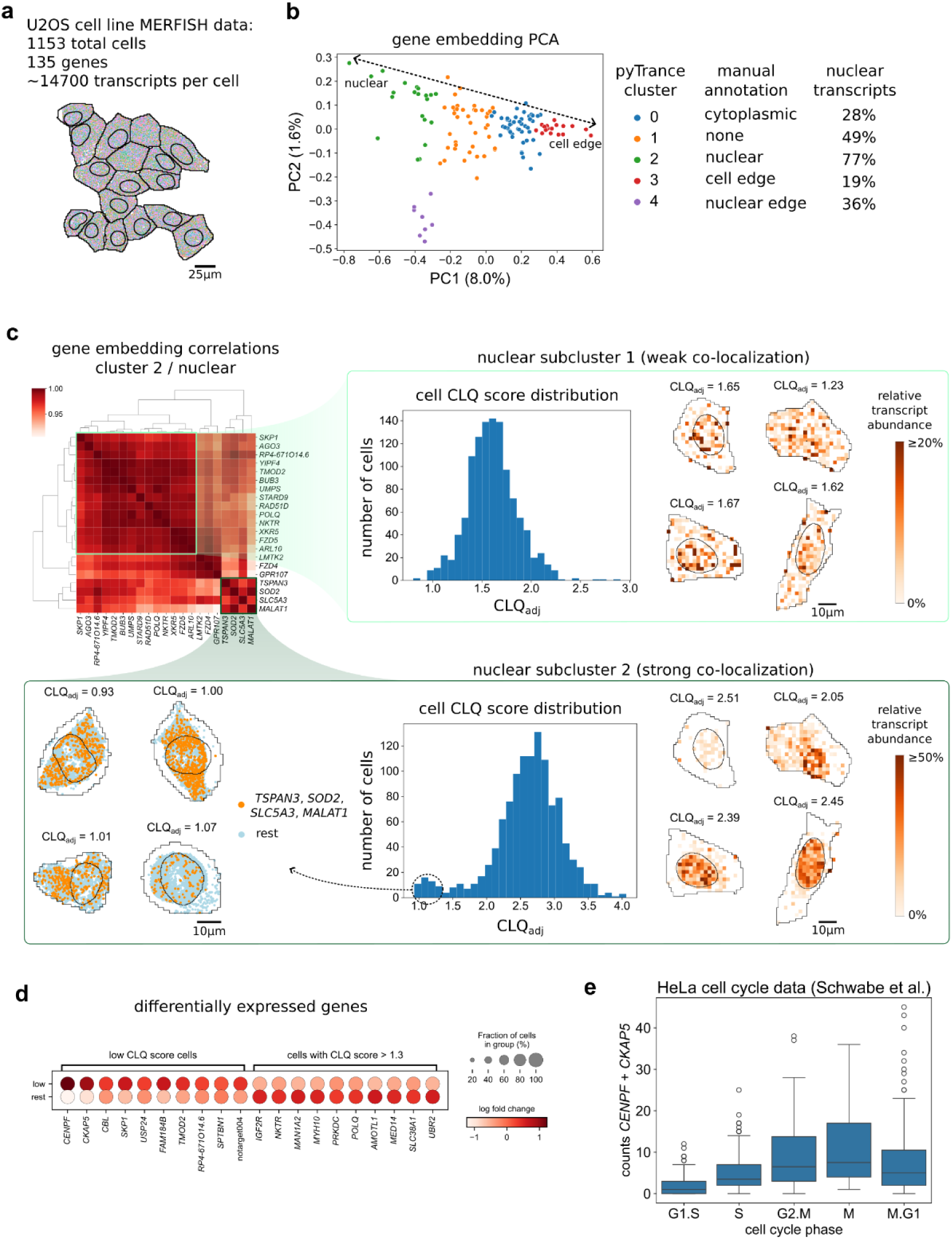
Dividing cells are identified by their distinct RNA localization. **a**, Overview of the used dataset. Image shows an example batch of cells, detected transcripts are shown as single dots colored by their corresponding genes. **b**, PCA of the gene embeddings computed with pyTrance colored by the clusters obtained from the embeddings space. **c**, Pairwise Pearson correlation matrix of gene embeddings of cluster 2 (top left). CLQ score distributions over all cells (middle) and relative transcript abundance histograms for randomly sampled cells (right) are shown for two highlighted gene subclusters. Single-transcript distributions of cells with low CLQ score for subcluster 2 on bottom left. **d**, Differentially expressed genes between cells with low (≤ 1.3) and high (>1.3) CLQ score of subcluster 2 genes. **e**, Summed CENPF and CKAP5 counts in an external HeLa cell dataset across cell cycle phases.

We clustered the gene embeddings computed with pyTrance and identified five clusters of co-localizing genes: four clusters corresponding to different cell compartments and one cluster containing genes following no specific localization pattern (**Fig. 4b**, **Extended Data Fig. 5a-d**). Our manual annotations are supported by the percentage of nuclear transcripts per gene cluster, which is highest for cluster 2 (“nuclear”) and lowest for cluster 3 (“cell edge”). We noticed that the CLQ scores computed for each cluster indicate a random localization in many cells of the “cytoplasm” and “cell edge” clusters (**Extended Data Fig. 5c**). This is most likely due to the radius-based graph construction, which does not reflect the ring-shaped localization to the cytoplasm and cell edge very well. Nevertheless, the patterns were detected by the GNN using the same graph, suggesting it is primarily a limitation of quantification using the CLQ score.

Upon further investigation of the nuclear gene set, we could identify two subsets with different co-localization strengths (**Fig. 5c**). For the gene set with strong co-localization (*TSPAN3*, *SOD2*, *SLC5A3*, *MALAT1*), we noticed a small outlier group of 45 cells, which had a CLQ score less than 1.3. The spatial transcript distributions of the respective genes confirmed the lack of co-localization indicated by the low CLQ score and further showed a common pattern of two separated transcript populations. Since many of these cells also lack a nuclear mask, we speculated that they in fact were dividing when the experiment was done (**Extended Data Fig. 5e**). This hypothesis was supported by their gene expression profiles, showing an upregulation of *CENPF* and *CKAP5*, two genes involved in chromosome segregation and spindle formation, respectively (**Fig. 4d**) (Gergely *et al*. 2003; Liao *et al*. 1995). Conventional single-cell clustering based on gene expression returned a cluster with similar marker genes, including *CENPF* and *CKAP5*, but with many cells exhibiting a strong localization for the nuclear gene set, suggesting no ongoing division (**Extended Data Fig. 5f-h**). Hence, we hypothesized that these cells have already accumulated *CENPF* and *CKAP5* RNA in preparation for division or have not yet degraded it after, making them hard to distinguish from the actually dividing cells based on just their transcriptional profiles. Indeed, in a scRNA-seq HeLa dataset annotated by cell cycle phase, we found RNA counts of *CENPF* and *CKAP5* to be very similar in G2.M, M, and M.G1 phases, supporting our hypothesis (**Fig. 4e**) (Schwabe *et al*. 2020). This underlines the benefit pyTrance provides by extending ST analysis beyond RNA abundance to subcellular localization.

**Fig. 5:**
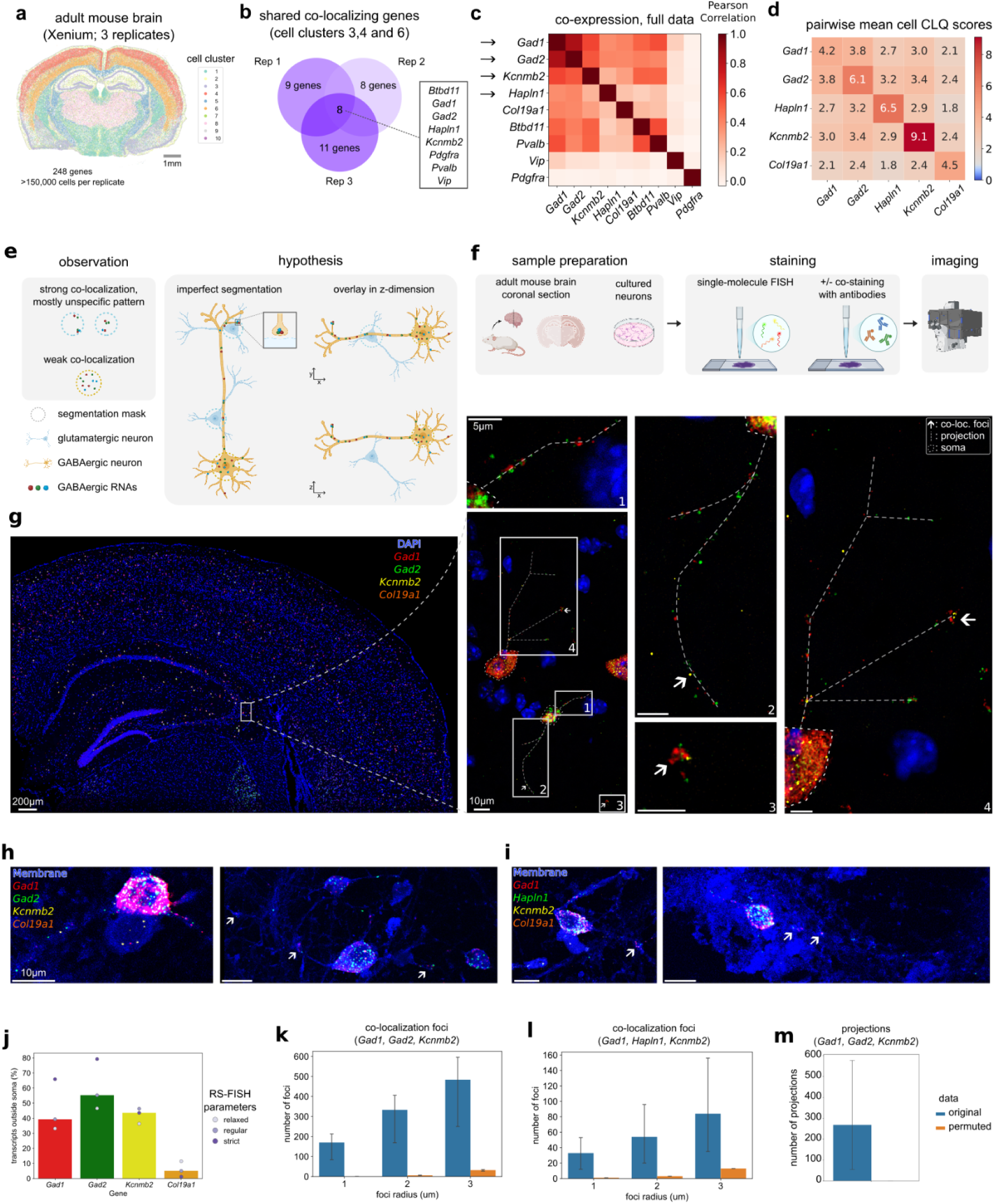
pyTrance identifies RNAs co-localizing in neuronal projections. **a**, Overview of the used dataset. Shown is the single-cell clustering on one replicate. **b**, Agreement of one pyTrance cluster between the three replicates. **c**, Co-expression of the pyTrance cluster shared between replicates computed on the full dataset. *Col19a1* is included as negative control regarding co-localization. Arrows indicate the candidates selected for further analysis and experimental validation. **d**, Pairwise CLQ score between the selected candidates. Shown is the mean across analyzed cells (cell clusters 3, 4, 6, all replicates). **e**, Overview of the observed co-localization (left) and the hypothesis that localization in projection and technical artifacts lead to the observed co-localization in different cell types. **f**, Overview of the performed validation experiments. **g**, Full smFISH image of one mouse brain section (left), close-up of a selected field of view and four regions of interest (ROI) therein (right). Arrows indicate foci where the three included candidate RNAs co-localize, straight dashed lines indicate neuronal projections based on the RNA molecule arrangement, round dashed lines outline the somas the projections belong to. Same scale bar size for all ROIs. **h**, **i**, Example images of cultured neurons showing localization of RNAs to neuronal projections. Arrows indicate RNA aggregates in potential synapses. Same scale bar size for all images. **j**, Quantification of RNA localization based on three different parameter settings for molecule detection in brain section 1. **k**, **l**, Quantification of the number of co-localizations of three of the RNAs for different radii. **k**, brain section 1, **l**, brain section 1. **m**, Quantification of the number of projections in brain section 1. For **k**, **l**, **m**, error bars indicate values for different RS-FISH parameters settings (original) and their results after random permutations (permuted)

We compared clusters obtained with pyTrance to the results from Bento’s RNAflux and RNAforest, as well as the consensus annotation the authors performed for the localization of each gene individually (Methods) (Mah *et al*. 2024). As in the simulated data, grouping genes based on their highest fluxmap values did not give meaningful results for RNAflux, hence, we again clustered them based on all fluxmap dimensions (**Extended Data Fig. 6a, b**). Contrary to our previous results, we found the gene clusters obtained with RNAflux and pyTrance to largely agree, whereas the RNAforest predictions and consensus annotation were more distinct (**Fig. 4b**, **Extended Data Fig. 6b-d**). Nevertheless, there was a strong agreement between pyTrance, consensus annotation, and RNAflux on the genes of pyTrance cluster 4 (“nuclear edge”). However, the additional genes in the consensus annotation and RNAflux showed no particular (co-)localization and were thus mislabeled/misclustered (**Extended Data Fig. 6e-g**). The genes of pyTrance cluster 4, on the other hand, showed a strong co-localization at the nuclear edge (**Extended Data Fig. 5d**). We observed a similar mismatch for the genes of pyTrance cluster 3 (“cell edge”) and the corresponding RNAflux cluster and consensus annotation (**Extended Data Fig. 5c, 6i-k**).

Taken together, pyTrance detected RNA co-localization in different compartments of immortalized cancer cells and revealed a subset of cells undergoing division, which could not be identified based on RNA abundance only, highlighting the added value of quantifying co-localization.

### pyTrance identifies a set of RNAs localizing in projections of mouse GABAergic neurons

Lastly, we applied pyTrance to an adult mouse brain dataset generated with the 10x Xenium technology consisting of three consecutive tissue slices (10x Genomics, 2023). We used the cell clustering provided with the data, which corresponds to the main brain regions (clusters 3, 4, 6, and 8: hippocampus, outer cortex, inner cortex, and thalamus, respectively) as well as different cell types defined by their expressed markers (GABAergic neurons (cluster 1), astrocytes (cluster 2), oligodendrocytes (cluster 5), endothelial cells (cluster 7), and microglia (cluster 10)) (**Fig. 5a**, **Extended Data Fig. 7a**, Methods). Since RNA localization is known to be an important mechanism of post-transcriptional regulation in neurons (Holt *et al*. 2019), we focused our analysis on the brain regions with higher enrichment of neuronal cells: cortex (clusters 4 and 6) and hippocampus (cluster 3). We ran the pyTrance GNN and subsequent gene embedding clustering for each of the three replicates. To avoid having differential gene expression as a confounder, we only included one cell cluster per run.

pyTrance identified (sub)clusters of co-localizing RNAs that share common cellular functions (**Extended Data Fig. 8a-d**). For example, (1) one gene set contains the well-known neuronal projection-localized RNA of *Arc* (Farris *et al*. 2014; Steward & Worley 2001), which pyTrance predicted to co-localize with several genes encoding for synaptic proteins (*Arhgap12*, *Grik3*, *Lypd6*, *Orai2*, *Cplx3*, *Syt6*, *Unc13c*, *Nxph3*, and *Gng12*), suggesting this gene set corresponds to dendritic localization (**Extended Data Fig. 8n, r**). This is further supported by their low abundance in the nuclei. (2) Additionally, a gene set showing perinuclear localization contains transcripts of *Cd24a*, *Cntn6*, *Col19a1*, *Col6a1*, *Ndst3*, *Pcsk5*, and *Trpc4* (**Extended Data Fig. 8h, l**). All of these genes correspond to perineuronal, extracellular matrix, or cellular membranes’ components, which are typically locally translated at the endoplasmic reticulum. This may explain the predicted co-localization and observed pattern. Out of all predicted RNA co-localization clusters, one stands out in terms of its RNA embeddings (**Extended Data Fig. 7b**). It comprises mostly markers for GABAergic neurons and is largely shared across the three replicates and brain regions (**Fig. 5b**, **Extended Data Fig. 7c, d**). Contrary to the examples above, these RNAs show no localization to a specific cell compartment but instead form small aggregates scattered within the cells (**Extended Data Fig. 7e**). We chose to validate the predicted co-localization of these RNAs due the unusual pattern, for which we narrowed down the gene set to four candidates: *Gad1*, *Gad2*, *Kcnmb2,* and *Hapln1*, as well as *Col19a1* as a negative control, which is co-expressed but not predicted to co-localize with the other four RNAs based on the embedding clustering (**Fig. 5c**). The candidates were selected from the initial gene set based on co-expression and cell type specificity (Methods). Computing the CLQ score for the four candidates as a group statistically confirms their co-localization (**Extended Data Fig. 7f**). Scores for RNA pairs were lower for pairs including *Col19a1*, supporting the prediction of lack of co-localization with the four candidates (**Fig. 5d**).

Even though these genes are markers for GABAergic neurons, we often found high expression of *Slc17a7* (vGluT1), a marker for glutamatergic neurons, in the same cells (**Extended Data Fig. 7g, h**). Furthermore, CLQ scores are high in particular for cells with high abundance of *Slc17a7* but low abundance of GABAergic markers (**Extended Data Fig. 7i**). Additionally, based on the initial cell clusterings on the full sections and the respective cluster markers, GABAergic neurons form a separate cluster not included in our analysis (**Extended Data Fig. 7a, j**). Thus, we hypothesize that the selected genes (*Gad1, Gad2, Kcnmb2, Hapln1)* are transcribed in the nuclei of GABAergic neurons, but their corresponding RNA molecules co-localize in neuronal projections beyond the cell body, potentially even within axosomatic terminals at glutamatergic neurons (**Fig. 5e**). Because the cell segmentation relies on DAPI staining, transcripts located in these distal neuronal projections are misassigned to neighbouring cells during data preprocessing due to imperfect segmentation and signal overlap along the z-dimension (Mitchel *et al*. 2026; Tiesmeyer *et al*. 2026). Together, these complexities create the misleading appearance of mixed GABAergic and glutamatergic RNAs within the same cell. To test this hypothesis directly, we performed 4-plex smFISH on adult mouse brain sections and cultured cortical neurons (**Fig. 5f**, Methods).

In the tissue sections we observed many examples of co-localization foci outside of somas, as well as patterns resembling neuronal projections (**Fig. 5g**, **Extended Data Fig. 9a**). Some of the foci are located at the boundaries of cells that do not show any other signal for the tested RNAs, indicating that these might actually be axosomatic terminals (**Extended Data Fig. 9b**). We observed the same patterns in cultured neurons, and by using a membrane stain, we could confirm that the RNA molecules are actually in projections and not the extracellular space (**Fig. 5h, i**, **Extended Data Fig. 9c**). To obtain coordinates of individual molecules and further quantify our observations on the tissue sections, we used RS-FISH (Methods). For the four candidates (*Gad1*, *Gad2*, *Kcnmb2*, *Hapln1),* we found 40-50% of the detected molecules to be localized outside of the soma, compared to ≤10% for the negative control (*Col19a1*) (**Fig. 5j**, **Extended Data Fig. 9d**). In one section we detected hundreds of co-localization foci and projection-like localization patterns (**Fig. 5k, m**). Although the absolute numbers were lower in the second section, this difference can largely be explained by the lower expression of *Hapln1* compared to *Gad2* (∼5.5 times fewer molecules detected, depending on the RS-FISH parameters) and the smaller imaged area (14084 vs 18149 detected nuclei) (**Fig. 5k, l**). The congruence of an approximately 6-fold difference between the sections across parameter settings supports a similar localization of all four RNAs in both sections and highlights the robustness of the analysis. Importantly, when permuting the molecule coordinates or including *Col19a1* the number of detected foci and projections dropped to almost zero, showing that the patterns observed cannot be explained by random fluctuations and confirming the lack of co-localization with *Col19a1* (**Fig. 5k-m**, **Extended Data Fig. 9e, f**).

To test if some of the localized RNAs could also be locally translated, we selected *Gad1* due to its high expression and the known constitutive function of its protein (Gad67) throughout the cell. We co-stained *Gad1* RNA and protein in mouse brain tissue (**Fig. 5f**, Methods). Focusing on the RNA molecules more distant from nuclei, we indeed find a strong co-localization with the protein signal, supporting our hypothesis of local translation in projections (**Extended Data Fig. 9g-i**).

## Discussion

We introduced pyTrance, a three-stage computational workflow for the systematic investigation of RNA co-localization in iST data. Its core functionality is the self-supervised learning of latent gene embeddings using a GNN based on the spatial proximity of transcripts. The embeddings are clustered to obtain co-localizing candidates, which are statistically validated using an adjusted version of the CLQ.

Using simulated data, we demonstrated pyTrance’s ability to successfully infer co-localization of RNA groups from individual localizations to the same cell compartments. Our introduced permutation-based adjustment of the CLQ results in an interpretable statistical measure of co-localization, which gives users an intuitive understanding for high or low values and allows a more reliable comparison of results for different sets of genes or cells.

Benchmarking against existing methods showed that pyTrance performs advantageously in terms of scalability and unbiased detection of co-localizing RNAs. Transferring the embedding and clustering-approach employed by pyTrance also lead to an improvement of results for RNAflux, presumably because it makes use of the information encoded in all latent dimensions instead of just a single one and is therefore better suited to deal with genes with weak or variable localization across cells. We note that the field of subcellular analysis is broad, and methods are designed with different applications in mind. Therefore, not all methods are directly comparable but could instead be used complementarily to validate one method’s predictions with an orthogonal approach or to further categorize co-localization groups.

We demonstrated the applicability of pyTrance to real data in two examples. In immortalized cancer cell line data, the gene clusters again corresponded to localization patterns in the main cell compartments. Follow-up analysis based on the CLQ score allowed us to identify dividing cells, which we failed to do using only RNA abundance levels, highlighting the additional value of analyzing RNA localization. The big differences between the provided consensus annotation and the consistent patterns we found using pyTrance highlight the difficulty of manually labeling single genes by their localization. This problem is further amplified in tissue data, which usually have much higher sparsity and expression differences between cells. Furthermore, it underlines the advantage of self-supervised methods that do not rely on, potentially faulty, manual annotations. Instead, the pyTrance clusters could be used as a starting point for an annotation and manually refined based on subclustering and statistical metrics like the CLQ score.

In a mouse brain dataset, we discovered a set of GABAergic neuron markers (*Gad1*, *Gad2*, *Hapln1*, *Kcnmb2*) localizing to projections and, presumably, synapses based on our analysis done with pyTrance. Multiple previous large-scale studies show no agreement regarding the localization of these four RNAs (Cajigas *et al*. 2012; Mendonsa *et al*. 2023; Middleton *et al*. 2019; Perez *et al*. 2021; Poon *et al*. 2006). One explanation could be a cell state-dependent or subtype-specific localization, as in our data we only observe the localization for a subset of GABAergic neurons. Importantly, the distinct cellular roles of the GAD isoforms themselves may further contribute to these discrepancies: while Gad67 (*Gad1*) is broadly distributed throughout the cell and provides GABA for processes beyond neurotransmission, Gad65 (*Gad2*) is preferentially localized to nerve terminals where it supports synaptic GABA release (Kaufman *et al*. 1991). These isoform-specific functions are regulated by different cellular mechanisms and imply distinct spatial requirements for mRNA localization and local translation. Thus, activity-dependent regulation, neuronal subtype composition, or differences in synaptic maturation states across datasets could shift the balance between somatic and projection localization, potentially explaining the inconsistent observations reported previously in literature (Buddhala *et al*. 2009; Kanaani *et al*. 2015).

The strong RNA-protein co-localization we observed for *Gad1* is consistent with the possibility of local translation, which, to the best of our knowledge, has not been reported before and presents a potentially new regulatory mechanism of local GABA synthesis that allows cells to quickly and efficiently synthesize new proteins upon physiological requirements. At the same time, our findings of RNA misassignment in the Xenium data further underline the inherent challenges of analyzing ST data on a single-cell level, particularly for cell types with complex morphologies like neurons, where transcript contamination from nearby structures can lead to erroneous cell type assignment and downstream analysis (Mitchel *et al*. 2026; Tiesmeyer *et al*. 2026).

The high CLQ scores observed for RNA pairs including *Col19a1* computed on the Xenium data appear to contradict the distinct localization patterns we observed in our smFISH validation experiments (**Fig. 5d**, **j**). We can attribute this discrepancy, again, to the misassignment of co-expressed transcripts discussed above, however, in this case affecting somal transcripts. This leads to high co-localization scores for metrics computed on individual cells. On the other hand, the initial embedding-based clustering of pyTrance, incorporating information across cells, correctly predicted the co-localization confirmed by our validation experiments, highlighting a key advantage of using a more complex model for hypothesis generation first.

### Limitations

The types of localization patterns pyTrance can detect depend on the way the transcript neighborhood graph is constructed. Currently, only a radius-based construction is implemented, which is not very suitable for ring-shaped patterns, as we have already observed, especially for the cell edge-localized RNAs. Different construction approaches could help to identify more diverse patterns or cluster RNAs more confidently.

The GNN architecture used here is a simple single-layer GCN. When experimenting with changes in the architecture (e.g., number of layers and latent dimensions or types of layers), we did not observe significant changes in the results, however, a more careful model selection could still uncover more nuanced co-localization (sub)clusters. Especially including distances and angles in the neighborhood graph using geometric learning might be better suited to deal with the given spatial data and could further alleviate limitations in the graph construction discussed above (Bronstein *et al*. 2021). However, more complex models also require longer training and potentially more data and finding the right model for a given dataset is a non-trivial task that also depends on the computational resources available.

Currently, we explore co-localization clusters using the pairwise gene embedding correlations and visual assessment of example cells. Applying explainability methods to the GNN output could extract more information contained in the embeddings, leading to a better understanding of the similarities and differences between clusters. Furthermore, investigating the single transcript embeddings before aggregation could reveal additional aspects of localization, e.g., general RNA polarity in a cell.

Furthermore, in this work we only showed examples of iST datasets, but our approach can also be adapted to subcellular sequencing-based ST data, such as Open-ST ((Chen *et al*. 2022; Schott *et al*. 2024)) or Stereo-seq (Chen *et al*. 2022; Schott *et al*. 2024). Primarily, the permutations to train the GNN and compute the CLQ score have to happen on a transcript, instead of a capture spot level. We did implement these changes, but the GNN failed to identify any true co-localization, which we suspect is mostly for two reasons. Firstly, the sections for sequencing-based methods are usually 10 μm thick and, unlike with most iST methods, are processed in one piece, which reduces the effective resolution. Secondly, the coverage of the entire polyadenylated transcriptome comes at the cost of reduced sensitivity and increased data complexity, which makes it significantly harder to find co-localizations. The second point also applies to iST methods with panels of several thousand RNAs, where we also failed to obtain meaningful results. A more complex model paired with the increasingly large datasets being generated might be able to address this problem. Similarly, we also envision an adaptation to raw high-plex image data to be possible, including spatial proteomics technologies like MIBI-TOF (Keren *et al*. 2019) or CODEX (Black *et al*. 2021). However, this would require more adaptations because color channels have intensity values instead of transcript counts. Finally, training the GNN on multiple cell types leads to RNA clusters corresponding only to the respective markers, which limits its application to a single cell type per run. This could be addressed by including cell type information as a covariate, which would allow training the model just once for a full data set.

In conclusion, our pyTrance framework extends the currently still small set of available analysis methods for detecting RNA co-localization patterns from ST data and, as we have shown, it can lead to interesting, new, and biologically meaningful discoveries. pyTrance is available as an open-source package at https://github.com/rajewsky-lab/pytrance/.

## Methods

### pyTrance framework

#### GNN

The GNN input consists of an expression matrix and a transcript neighborhood graph per cell. The expression matrix naturally follows a one-hot encoded format such that each row has exactly a single 1 entry with the other entries being 0, where each row stands for a transcript and each column for a gene. The neighborhood graph is computed for each cell individually, such that transcripts of different cells have no connection. Extracellular transcripts are excluded. To construct the graph, the radius_neighbors_graph function from sklearn (v1.5.2) was applied to the transcript’s spatial coordinates, with the radius parameter depending on the dataset.

As a training scheme the self-supervised Deep Graph Infomax DGI framework is applied, using the implementation of the corresponding GitHub repository (https://github.com/PetarV-/DGI) (Veličković *et al*. 2018). The permuted input is constructed by shuffling gene labels within a cell while the neighborhood graph and transcript coordinates stay fixed. Only a single GCN layer is used such that only information of a transcript’s direct spatial neighbors is taken into account. The model is trained using the Adam optimizer with default parameters (Kingma & Ba 2014). The model output is a latent embedding for each transcript, where the number of latent dimensions depends on the dataset.

#### Embedding analysis

Gene embeddings are obtained by computing the mean of all corresponding transcript embeddings for each gene. For the gene embedding clustering, we use the Leiden clustering implementation of scanpy (v1.10.3) (Wolf *et al*. 2018). Principal Component Analysis on the gene embeddings was performed using the decomposition.PCA.fit_transform function of sklearn (v1.5.2). Gene embedding correlations were computed using the DataFrame.corr function of pandas (v.2.2.3) and correlation heatmap structures are determined using agglomerative clustering as implemented in sklearn.

When working with unsupervised clustering algorithms, the exact number of clusters to expect is usually unknown, and boundaries between clusters can be blurry. Therefore, we recommend exploring the initial clusters and adjusting the clustering parameters if necessary.

#### Cell CLQ score

The raw CLQ is defined as:

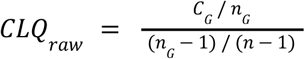

Where *C* _*G*_ is the number of connections between transcripts of the gene set *G* in the neighborhood graph, *n* _*G*_ is the number of transcripts of the gene set *G* and *n* is the total number of transcripts in a given cell. This differs from the original CLQ definition of Leslie & Kronenfeld in that all connections within the gene set are counted, instead of only nearest neighbors (Leslie & Kronenfeld 2011).

To compare the CLQ to a random baseline, the gene labels are shuffled repeatedly and *CLQ* _*CLQ*_ is recomputed after each permutation, such that *CLQ* _*perm, i*_ defines the *CLQ* _*CLQ*_ value on permutation _*i*_ defined as of a given cell. The adjusted CLQ, here called cell CLQ score, is then

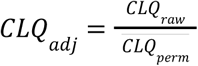

Where *CLQ* _*perm*_ is the mean over all *CLQ* _*perm, i*_.

For all results shown here the neighborhood graph of each cell is recomputed with the same parameters as used for the GNN input graphs. The number of permutations was set to 100. Only cells with at least 5 transcripts of the respective gene set were considered for cell CLQ score computation.

### Relative transcript abundance

Relative transcript abundances were computed using the histogram2d function of NumPy and define the percentage of transcripts from a given gene set out of the total transcripts within a spatial bin.

### Pattern terminology

To overcome ambiguous naming for localization patterns across literature, we used the following terms consistently in our work: ‘nuclear’, ‘nuclear edge’, ‘cytoplasm’ and ‘cell edge’ describing a predominant localization of the transcripts at the respective compartments, ‘foci’ describing an aggregation of transcripts in small hotspots independent of the cellular compartment and ‘random’ describing a random localization across the cell. These names deviate from some of the ones used originally for the datasets and methods used here.

### Bento RNAflux

Fluxmaps were computed using the tl.flux, tl.fluxmap, and tl.comp functions of the bento-tools package (v2.1.4). If not stated otherwise the functions were run with default parameter values.

### Bento RNAforest

RNAforest was run using the tl.lp and tl.lp_stats functions with default parameter values of the bento-tools package (v2.1.4). The gene labels were obtained by aggregating the binary predictions for each gene based on a majority vote across cells.

### Simulated data

#### Simulation

The data were generated using the simulate_localization_pattern function of the Sim-FISH package (v0.2.0). For a total of 5000 cells, the 317 nuclei and cell outlines provided by the package were used repeatedly such that each template was used 15-16 times. For each of the ‘foci’, ‘nuclear’, ‘cytoplasm’, ‘nuclear edge’ (called ‘perinuclear’ by Sim-FISH) and ‘cell edge’ (called ‘pericellular’ by Sim-FISH) localization patterns, 30 genes were simulated per cell as well as 300 genes with a ‘random’ localization pattern. The framework also includes a ‘protrusion’ pattern, which, however, is only available for half of the provided cell shapes and was therefore not used for the analysis here. For each pattern the respective genes were split equally into 3 groups with mean expressions per cell of 5, 10, and 15, respectively. The number of spots per gene was sampled for every cell individually using a normal distribution with a variance of 20 and a mean depending on the expression group. Negative sampled expression values were set to 0. The gene proportion parameter defining the pattern intensity was sampled for every gene using a mean of 0.5 and variance of 0.3, and clipped to a range of 0 to 1. The proportion was kept fixed over all cells for each gene.

#### Analysis

The neighborhood graph was constructed using a radius of 20 and a GCN layer with 32 latent dimensions was used. The gene embeddings were computed as described above and clustered using the Leiden clustering algorithm implemented in scanpy (v1.10.3). The resolution parameter was tuned manually to obtain 6 clusters, the same number as simulated localization patterns. The gene subset of cluster 2 was chosen manually based on within-group correlation and the branching of the hierarchical clustering tree. Cell CLQ scores were computed as described above. The example cells shown in Fig. 2F were chosen manually based on their respective raw and adjusted CLQs. For the random control group the same number of genes as in cluster 2 were sampled randomly from the genes with a ‘random’ localization pattern.

### Benchmark

#### Bento RNAflux

The number of fluxmaps was set to 6, the same number as simulated localization patterns.

#### Bento RNAcoloc

RNAcoloc was run using the tl.coloc_quotient and tl.colocation functions of the bento-tools package (v2.1.4). The number of ranks was set to 6, the same number as simulated localization patterns. Since the pairwise co-localization results cannot be compared to the clusters computed using pyTrance, RNAcoloc could not be included in the performance analysis.

#### InSTAnT

InSTAnT was run using its run_ProximalPairs and run_fsm functions with default parameter values and run_GlobalColocalization with a distance threshold of 20 and min_genecount of 10. run_fsm returned an empty table, a problem we were not able to fix even with the help of the InSTAnT developers. Since the pairwise co-localization results cannot be compared to the clusters computed using pyTrance, InSTAnt could not be included in the performance analysis.

#### FISHfactor

FISHfactor was run using its model.inference function with 6 factors. After 72h of computing it was terminated. Even when tweaking the model parameters, the runtime could not be reduced significantly, hence, FISHfactor was not considered for further analysis.

#### Sprawl

Sprawl was run using its scoring.iter_scores function for the four available metrics. After 72h of computing it was terminated and not considered for further analysis.

#### ELLA

ELLA was run for individual genes using the same parameters as in the mini demo the authors provide.

#### CellSP

CellSP was run using the ch.run_instant and ch.bicluster_instant functions. We tried different parameter settings, however, the resulting dataframe in the anndata object was always empty, so we could not apply CellSP to the InSTAnT or Sprawl output.

#### Compute hardware

All methods were run on an Intel Xeon Platinum 8280 CPU. pyTrance and FISHfactor, which support GPU usage, were trained on a Nvidia A100.

#### Analysis

For the performance analysis, the number of cells in the full data set was downsampled to 100, 500, 1000, 2500, or kept at 5000. Additionally, for genes that are not ‘random’ the proportion parameter was thresholded to a minimum of 0.1, 0.2, and 0.5 within the cell subsets, such that only genes with a localization intensity higher than the respective threshold are kept. For subsets with no intensity threshold, the total number of transcripts was downsampled to 10% and 50%. For each resulting dataset, the GNN was trained with the same hyperparameters as used for the full dataset (radius: 20, latent dimensions: 32), and gene embeddings were computed as described above. The gene embeddings were clustered using k-means and agglomerative clustering with 6 clusters and Leiden clustering with resolution 0.6 for subsets with intensity thresholds and 0.65 for subsets with downsampled transcript numbers. For each clustering the ARI and AMI scores were computed. Both scores were computed using the respective functions implemented in sklearn (v1.5.2). The Leiden resolution parameter was manually chosen such that the number of clusters was close to the number of simulated patterns for most subsets, without having to optimize the parameter for each case individually. For the full dataset we additionally computed the ARI score after each epoch of GNN training for pyTrance (**Extended Data Fig. 3a**).

To compute the results of Bento RNAflux and RNAforest, the cells first had to be shifted as they were simulated with respect to (0, 0) independent of each other, therefore, the coordinates overlap and otherwise cells would not be handled individually by the Bento io.prep function. To do so, molecule, cell, and nucleus boundaries were shifted in x incrementally for each cell such that they are ordered in a line without overlapping. Results for RNAflux and RNAforest were then computed as described above for each data subset. For faster processing, the number of workers was set to 3 and the resolution parameter of RNAflux to 0.1. Genes were either assigned to a single fluxmap based on the one with the highest value or by treating the fluxmaps as embeddings and clustering them as described above for pyTrance, but with a Leiden resolution parameter of 0.3. Scores for RNAflux clusters were computed as described above. Scores for RNAforest were computed by treating the class predictions as clusters.

#### U2OS analysis

The dataset is publicly available at https://doi.org/10.6084/m9.figshare.c.6564043. The neighborhood graph was constructed using a radius of 5 μm and a GCN layer with 16 latent dimensions was used. The gene embeddings were clustered using the Leiden clustering algorithm implemented in scanpy. Clusters were annotated based on visual inspection of transcript distribution in random cells. The percentage of nuclear transcripts per gene cluster was computed as the sum of respective transcripts localized in the nuclear mask divided by the total number of respective transcripts across all cells where a nuclear mask was available. The gene subsets of cluster 2 were chosen manually based on within-group correlation and the branching of the hierarchical clustering tree. Cell CLQ scores were computed as described above. The ‘consensus annotation’ is part of the public data for which the authors annotated genes for their localization in individual cells (see (Mah *et al*. 2024). To obtain a single label for each gene, a majority vote was performed on cell level annotations to determine which label to keep. If two labels had equal numbers of votes one was chosen randomly. The cells were clustered on a single-cell level using Leiden clustering. Differentially expressed genes were computed on normalized and log1p-transformed cells’ expression profiles using the tl.rank_genes_groups function implemented in scanpy.

The cell cycle phase annotations provided by Schwabe et al. were used for the HeLa cell data (Schwabe *et al*. 2020). The total CENPF and CKAP5 unique molecular identifier counts of the raw data were summed for each cell.

#### RNAflux & RNAforest

Results for both methods were recomputed. For RNAflux, the number of workers was set to 3 and the resolution parameter to 0.1 for faster processing. The number of fluxmaps was set to 5 to match the number of obtained clusters from pyTrance. Leiden clustering was used to cluster the genes based on their fluxmap embeddings.

#### Adult mouse brain analysis

The dataset is publicly available on the <u>10x Genomics website</u>. The provided k-means clustering with 10 clusters was used to identify cell types and tissue regions. Differential gene expression was computed using the scanpy tl.rank_genes_groups function and cell type clusters (2: astrocytes, 5: oligodendrocytes, 7: endothelial cells, 10: microglia) were annotated manually based on differentially expressed marker genes.

Because we wanted to investigate neuronal RNA localization, genes labeled markers for ‘Astrocytes’, ‘Endothelial cells’, ‘Oligodendrocytes’, ‘Microglia / perivascular macrophages’ in the supplemental gene list provided with the dataset were removed. The neighborhood graph was constructed using a radius of 2, including the z-dimension, and a GCN layer with 32 latent dimensions was used. Compared to the U2OS dataset, which has a similarly sized gene panel, the number of latent dimensions was increased to account for the higher complexity in tissue data. The gene embeddings were computed for clusters 3, 4, and 6 independently for each of the three replicates and clustered using the Leiden clustering algorithm implemented in scanpy. Co-expression was computed using the np.corrcoef function. To compute the pairwise CLQ scores, only cells with at least 2 transcripts of each of the two genes were considered. The scores are computed as described above for each gene pair and each cell in clusters 3, 4, and 6 of all three replicates. Because the pairwise score distributions are heavily skewed towards zero, the median score for most gene pairs is zero, therefore, the mean score was used to compare the results.

Because the RNAscope™ assay used for experimental validation only offers multiplexing of up to 4 targets, not all genes from the initial gene set could be validated. *Pdgfra* and *Vip* were excluded due to low co-expression. *Pvalb* was excluded because it is a marker for a specific type of interneurons and we wanted to keep the analysis as broad as possible. For *Btbd11* there was no probe available for RNAscope™.

### Validation experiments

#### RNAscope™ Multiplex Fluorescent RNA in situ hybridization

Single-molecule multiplex RNA fluorescence in situ hybridization was performed using the RNAscope Multiplex Fluorescent assay (RNAscope™ Multiplex Fluorescent Reagent Kit v2. Advanced Cell Diagnostics, ACD) according to the manufacturer’s protocol for fresh-frozen samples.

For tissue experiments, brains were dissected from postnatal day 60 C57BL/6N mice and embedded and then cryosectioned at 10 µm thickness. Sections were fixed in 4% paraformaldehyde at 4 °C, followed by dehydration through an ethanol series. Endogenous peroxidase activity was quenched using hydrogen peroxide treatment, and tissue permeabilization was achieved using the recommended protease pretreatment for fresh-frozen samples as specified by the manufacturer.

For cell culture experiments, primary mouse cortical neurons were prepared from newborns (P0) following standard protocols and maintained in culture for 14 days. Cells were fixed directly on coverslips using 4% PFA, followed by dehydration and pretreatment steps according to the RNAscope™ protocol for adherent cells.

RNAscope™ target probes were hybridized simultaneously using the multiplex assay protocol. The following mouse probes were used: *Col19a1* (channel C1, 690 nm), *Gad2* (channel C2, 520 nm), *Kcnmb2* (channel C3, 570 nm), *Gad1* (channel C4, 780 nm), and *Hapln1* (channel C2, 520 nm). Probe hybridization was performed at 40 °C in a humidified oven, followed by a series of signal amplification steps according to the RNAscope multiplex protocol. Fluorophores corresponding to each channel were developed sequentially. Nuclei were counterstained with DAPI, and ExoBrite™ 410/450 was used for membrane staining.

#### RNAscope™ Multiplex Fluorescent RNA in situ hybridization with Immunofluorescence

For combined RNA *in situ* hybridization and immunofluorescence of *Gad1* RNA, Gad1 protein, and vGLuT1 protein, the RNAscope assay was performed as described above. After signal amplification steps, the slides were subsequently processed for immunofluorescence. Primary antibodies for Gad1 (ab261166) and vGluT1 (Sysy 135318) proteins were incubated overnight and detection was performed using fluorophore-conjugated secondary antibodies, followed by Opal 780 fluorophore development and DAPI counterstain, according to the manufacturer’s recommendations.

#### Imaging

Selected fields of view were imaged on a Leica STELLARIS 8 FALCON STED confocal microscope using a 63X oil-immersion objective. Appropriate laser lines and detection windows were used for DAPI/Exobrite™ and RNAscope fluorophores. Imaging parameters were kept constant across samples. Built-in deconvolution software (*Leica LIGHTNING)* was used during acquisition to obtain higher quality images.

### Validation analysis

#### Segmentation

To segment soma, Cellpose (v4.0.7) was run on the channels for DAPI, *Gad1,* and *Gad2* (for section 2 only DAPI and *Gad1*). This segmented nuclei and, for cells expressing Gad, the somas very reliably. However, for cells with high Gad expression, we found the outer part of the soma to be missed in some cases. A separate Cellpose run on only the channels for *Gad1* and *Gad2* (for section 2 only *Gad1*) helped to detect those parts missed by the first run. The masks from both runs were merged and manually refined to remove last cells missed by Cellpose.

#### Spot detection

RNA molecule coordinates were obtained using the RS-FISH plugin of ImageJ. It was run with three different parameter settings for each of the two sections imaged. The parameters were chosen by visual assessment, such that ‘strict’ and ‘relaxed’ settings represent an upper and lower bound for the total numbers, where ‘strict’ includes only spots that are visually clearly identifiable as such and ‘relaxed’ includes more cases with lower certainty. The ‘regular’ setting serves as a compromise between the two. RS-FISH was run twice for each channel and parameter setting, once on the raw image and once on an image with pixels inside the soma masks set to a value of 0.

#### Quantification

The proportion of transcripts outside the soma was computed for each channel and RS-FISH parameter setting as follows:

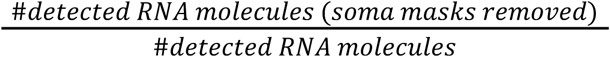

Co-localization foci were computed on the coordinates of the molecules detected with RS-FISH. The foci are defined as occurrences where at least one molecule of each RNA is present within a given radius.

Projections are defined as linear segments with a maximum length of 25 µm connecting two molecules, where each segment is required to have at least eight additional molecules within a maximum distance of 0.5 µm.

Permutations were computed by randomly shuffling molecule coordinates within the area spanned by the convex hull of the original coordinates. For each RS-FISH parameter setting, the coordinates were permuted 50 times. Co-localization foci and projections were recomputed on the coordinates of each permutation.

Segmented nuclei were counted by running ‘Analyze Particles’ in ImageJ on the binarized, merged and refined segmentation mask images.

## Acknowledgement

We thank Sandra Raimundo, Robert Zinzen, and the Systems Biology Imaging Technology Platform of MDC for the great support with imaging. Thanks to Anastasiya Boltengagen for the help with tissue cutting. We thank Marina Chekulaeva for discussions on neuronal RNA localization. Thank you to Zane Kliesmete, Tancredi Massimo Pentimalli, Florian Bartsch, and Joshua Günther for discussions on the manuscript. We thank all members of the Rajewsky lab for critical and helpful discussions. L.S. was financially supported by the International Research Training Group (IRTG) 2403 program of the Humboldt-Universität zu Berlin and the Helmholtz HPC Gateway Initiative. C.A.C-J. was financially supported by the HFMI Pilot Project: VirtualCell. N.R. thanks the Leibniz prize of the Deutsche Forschungsgemeinschaft (DFG) (grant number RA 838/5-1). Illustrations in Figure 5e were created with BioRender.

## Author contributions

L.S. and N.R. conceptualized and designed the project. L.S. implemented the method and performed the analysis. C.A.C-J. performed the *in situ* validation experiments and imaging with assistance from L.S.. N.K. and N.R. supervised the project. L.S., C.A.C-J., and NR wrote the manuscript. All authors edited and revised the manuscript.

## Extended Data Figures

**Extended Data Fig. 1:**
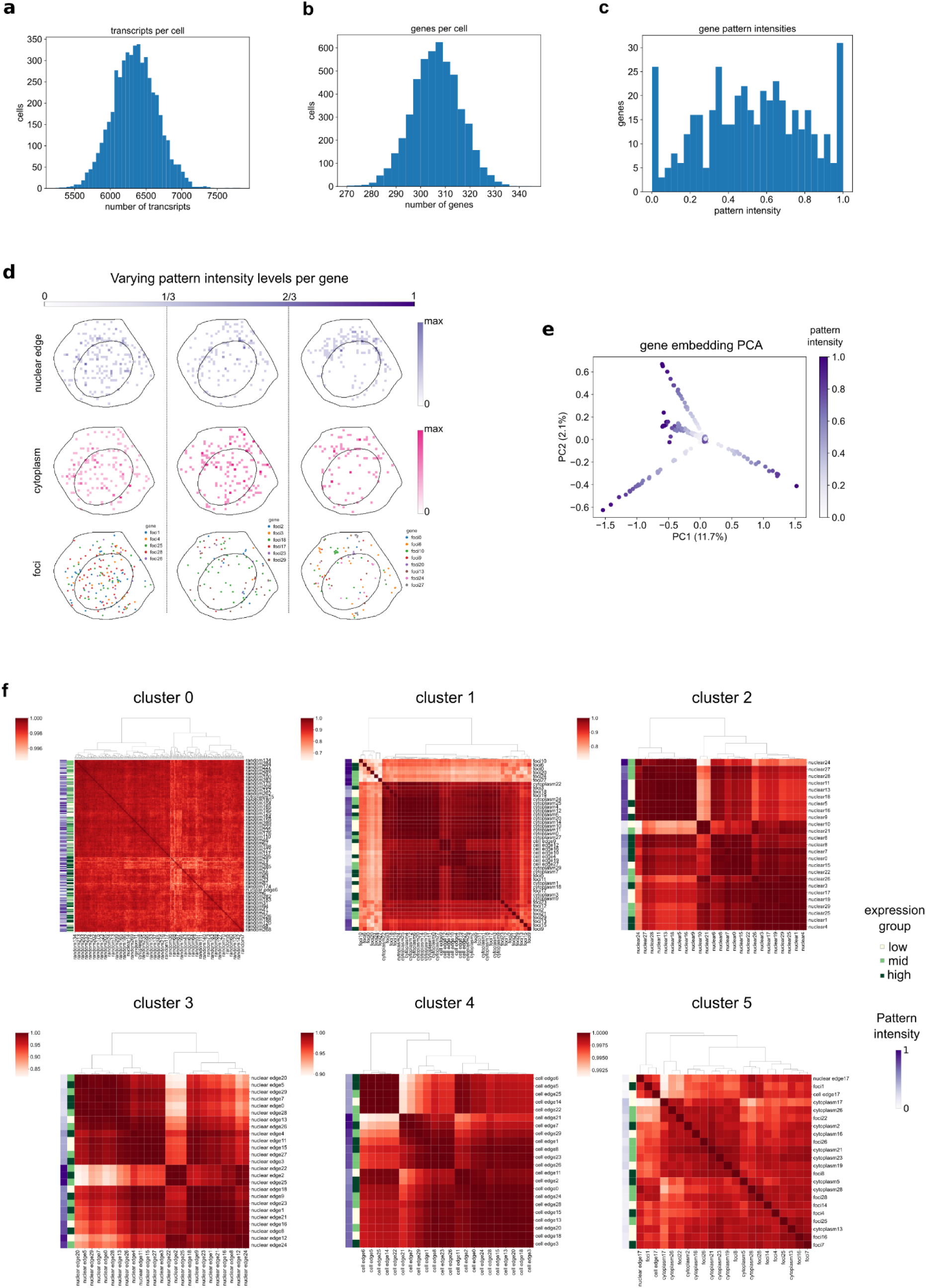
Sim-FISH data and corresponding pyTrance results. **a**, **b**, **c**, distributions of transcripts per cell (**a**), genes per cell (**b**) and gene pattern intensity (**c**) in the simulated data. **d**, Examples of different localization intensities for three patterns. ‘nuclear edge’ and ‘cytoplasm’ genes are shown cumulatively within each interval. **e**, PCA of the gene embeddings computed with pyTrance colored by the pattern intensity for each gene. **f**, Pairwise Pearson correlation matrices of gene embeddings of all pyTrance clusters. Additionally for each gene the simulated expression group and pattern intensity are shown.

**Extended Data Fig. 2:**
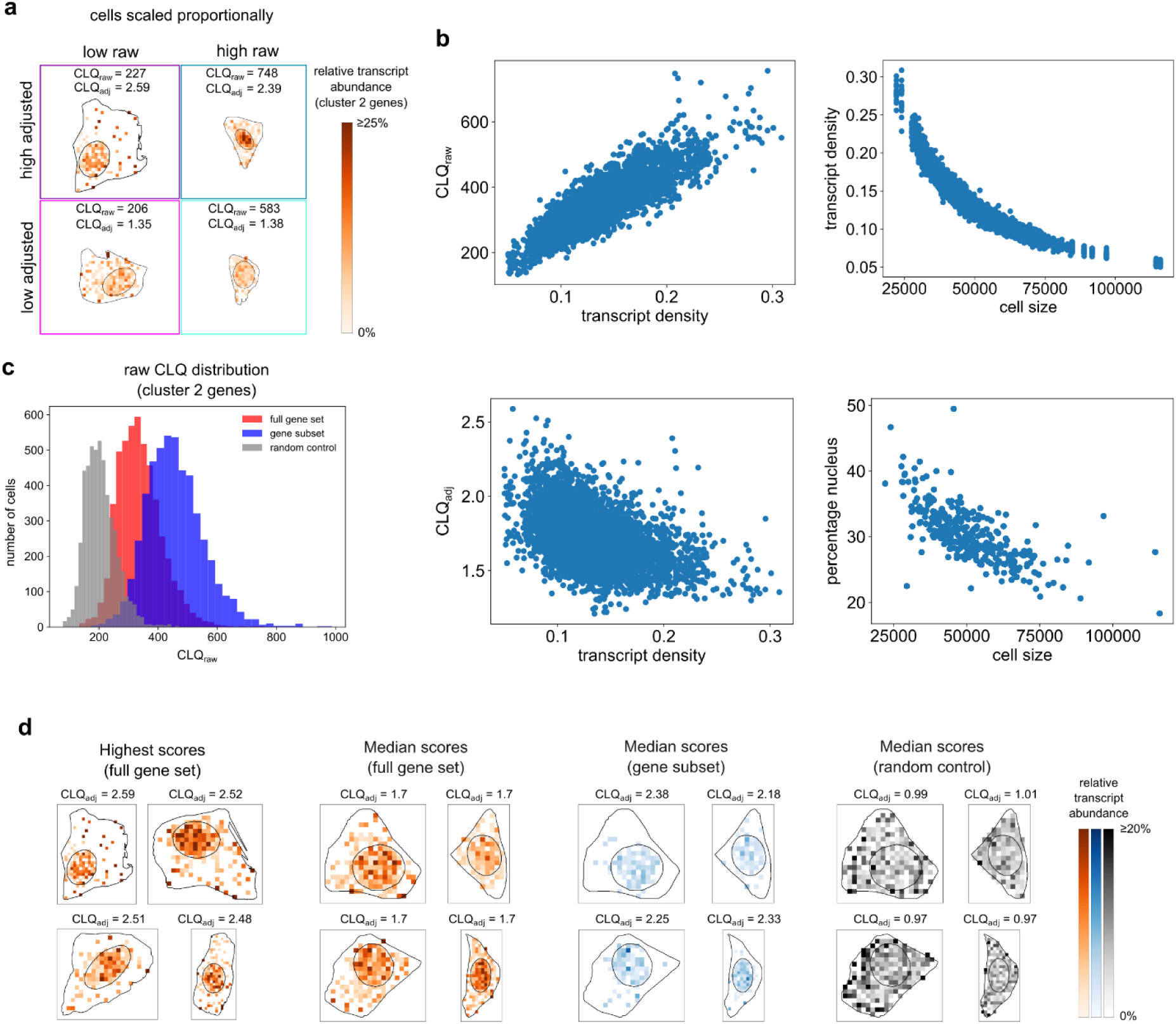
Comparison of raw and adjusted CLQ. **a**, Example cells with high/low raw and adjusted CLQs highlighted in Fig. 2e. Shown is the transcript count of the selected genes relative to the total counts per spatial bin (relative transcript abundance). Cells are scaled proportionally to their simulated size. **b**, Left: relationship between transcript density (number of transcripts / cell size) per cell and raw (top) and adjusted CLQ (bottom). Right: relationship between cell size and transcript density (top) and percentage of the cell area occupied by the nucleus (bottom). **c**, Distribution of raw cell CLQs for all cluster 2 genes (red), the subset with high embedding correlations highlighted in Fig. 2d (blue), and a set of genes with random localization (grey). **d**, Relative transcript abundance histograms of: all genes in cluster 2 for cells with highest (left), and median adjusted CLQ (middle-left), selected gene subset in same cells (middle-right), set of sampled genes with random localization in same cells (right).

**Extended Data Fig. 3:**
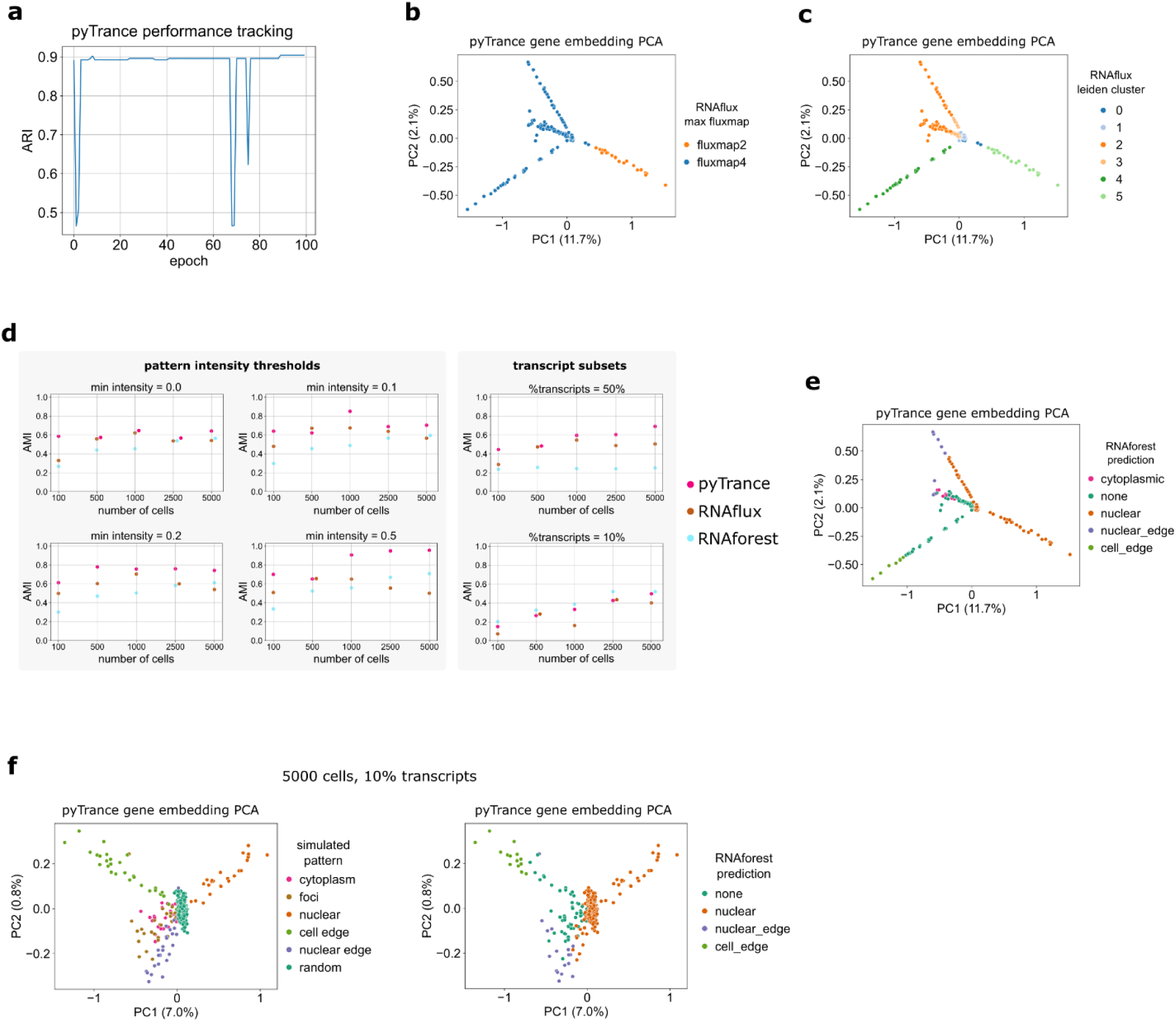
Extended performance benchmark of pyTrance, RNAflux and RNAforest. **a**, ARI tracked during training of the pyTrance GNN with the clusters recomputed after each epoch. **b**, **c**, PCA of the gene embeddings computed with pyTrance colored by the highest RNAflux fluxmap (**b**) and clusters obtained by clustering all fluxmap values per gene (**c**). **d**, The full simulated dataset was downsampled and filtered for genes with minimum simulated pattern intensities (left) or percentages of the total number of transcripts were sampled (right). AMI measures how well the computed clusters (pyTrance and RNAflux) or predicted labels (RNAforest) match the simulated patterns. **e**, PCA of pyTrance gene embeddings colored by RNAforest predictions. **f**, PCA of pyTrance gene embeddings colored by true pattern (left) and RNAforest predictions (right) for data where 10% of the total transcripts were sampled randomly.

**Extended Data Fig. 4:**
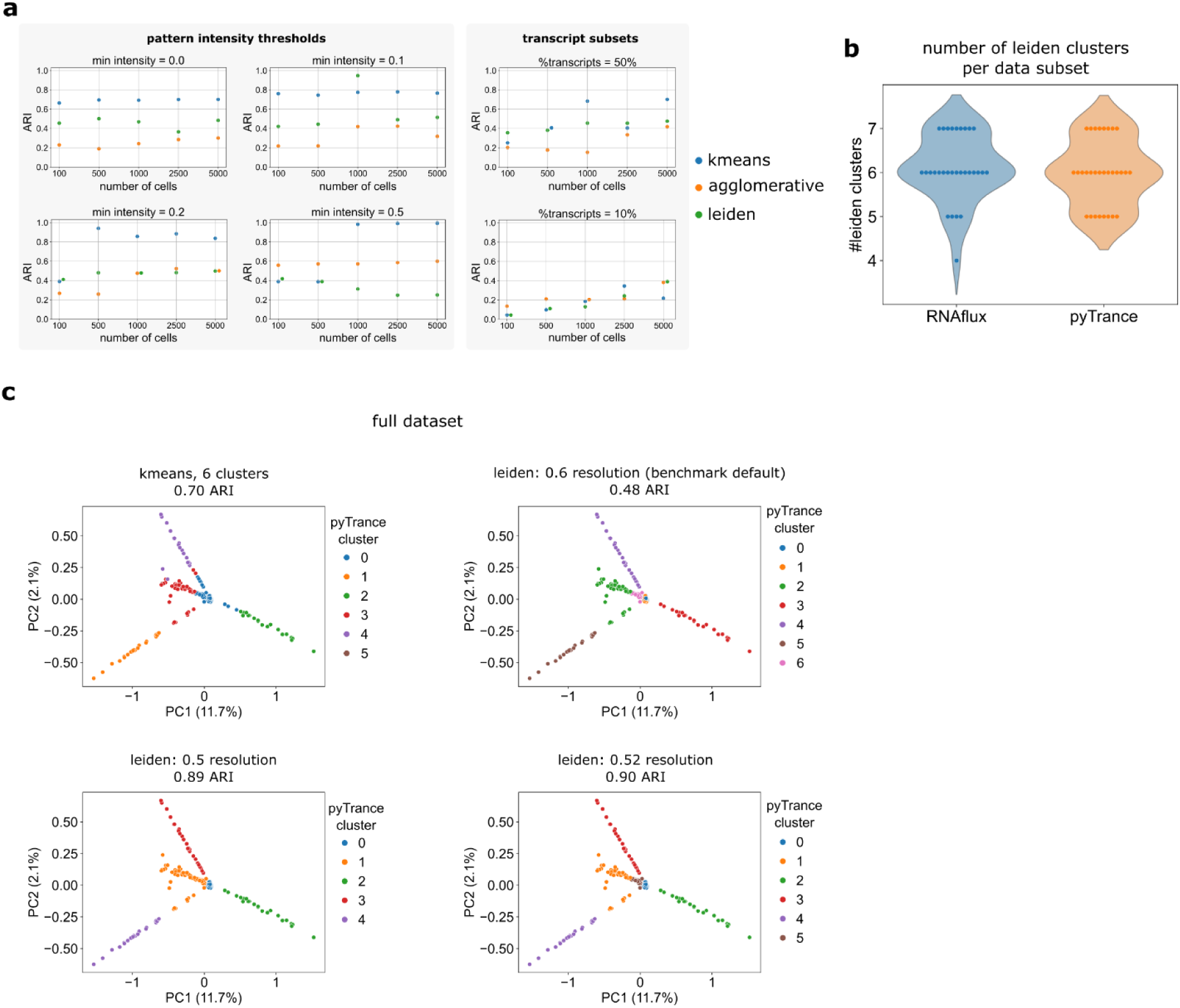
Comparison of gene embedding clusterings. **a**, Comparison of three clustering algorithms run on the pyTrance gene embeddings. The full simulated dataset was downsampled and filtered for genes with minimum simulated pattern intensities (left) or percentages of the total number of transcripts were sampled (right). ARI measures how well the computed clusters match the simulated patterns. **b**, Distributions of the number of obtained clusters for RNAflux and pyTrance by the pre-set Leiden parameters. **c**, Example of possible clustering improvement when manually fine-tuning the Leiden parameters.

**Extended Data Fig. 5:**
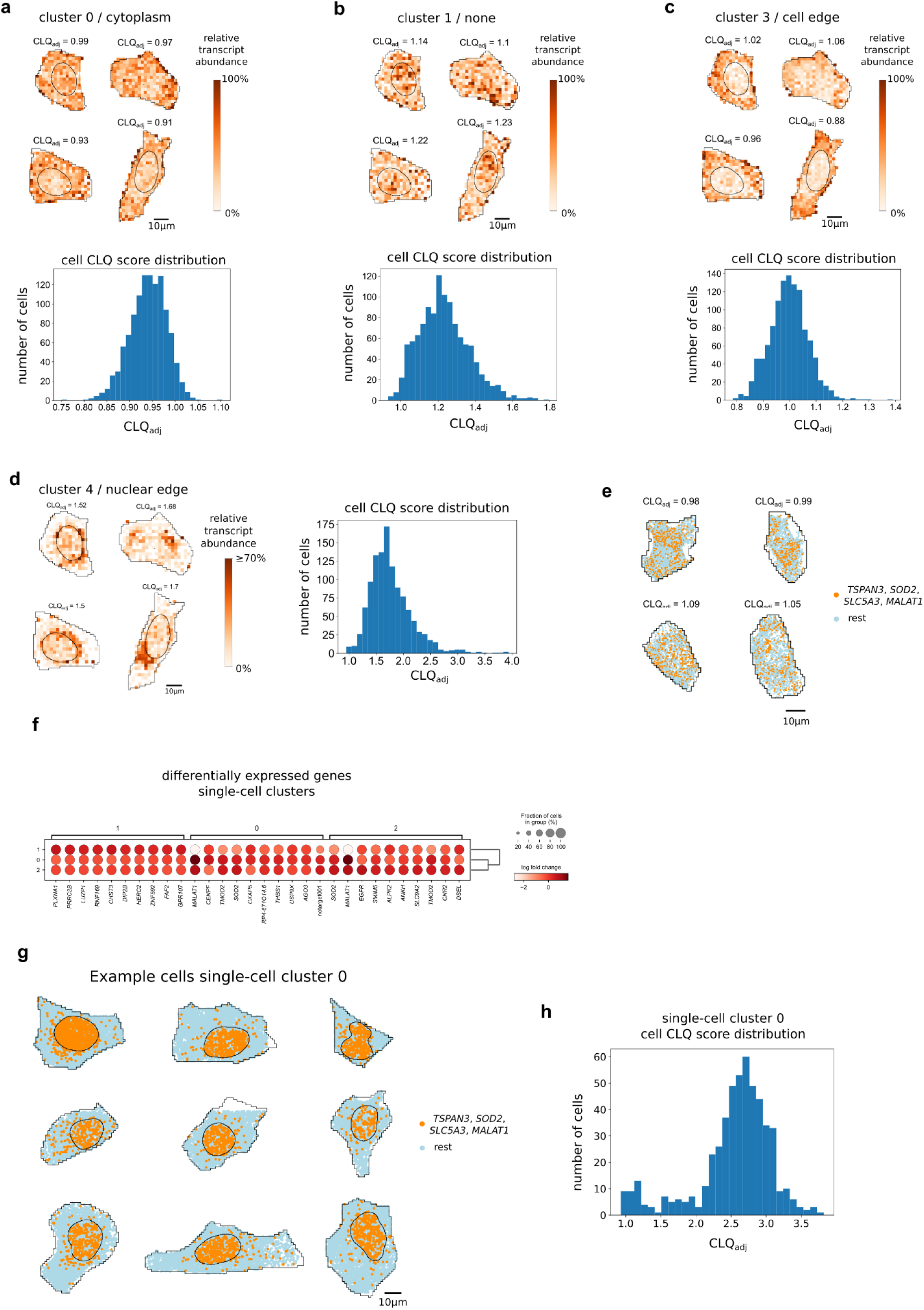
Extended pyTrance results on U2OS data. **a**-**d**, Relative transcript abundance histograms for randomly sampled cells and CLQ score distributions over all cells for four pyTrance clusters. **e**, Single-transcript distributions of RNAs with otherwise strong co-localization in the nucleus. Shown are example cells without a nuclear mask and no co-localization of the four RNAs. **f**, Differentially expressed genes of three clusters obtained by single-cell clustering. **g**, Single-transcript distributions of the same RNAs as in **e** in cells of single-cell cluster 0 where two mitotic markers (CENPF, CKAP5) are upregulated. **h**, Cell CLQ score distribution for the same RNAs in cells of single-cell cluster 0.

**Extended Data Fig. 6:**
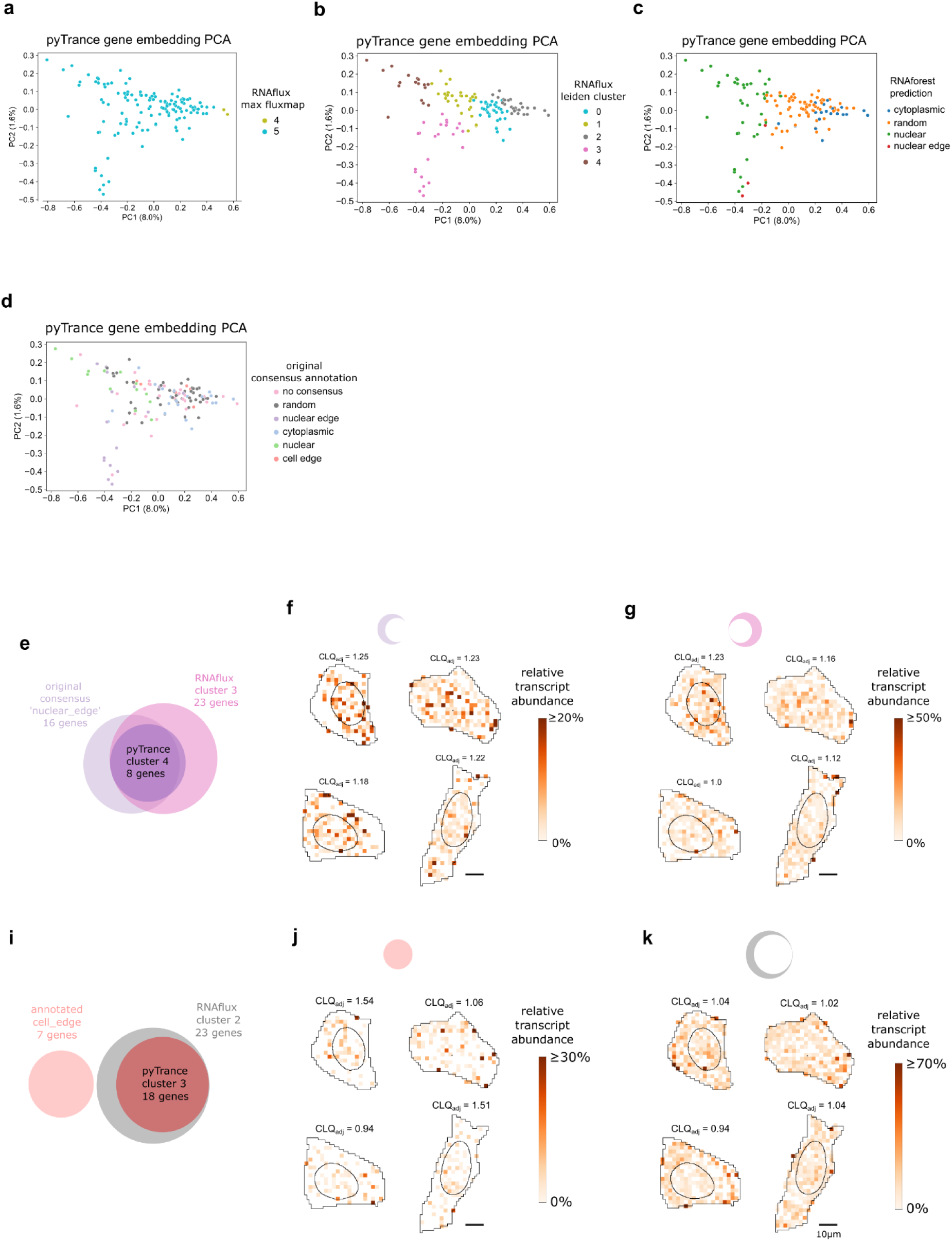
Comparison of pyTrance, RNAflux, RNAforest results and provided annotation on U2OS data. **a**-**d**, PCA of the gene embeddings computed with pyTrance colored by the highest RNAflux fluxmap (**a**), clusters obtained by clustering all fluxmap values per gene (**b**), RNAforest predictions (**c**), and the consensus manual annotation provided with the data (**d**). **e**-**k**, comparison of pyTrance clusters, RNAflux clusters and provided annotation for ‘nuclear edge’ RNAs (**e**-**g**) and ‘cell edge’ RNAs (**i**-**k**). Shown are the relative transcript abundance histograms for RNAs annotated (**f**, **j**) or clustered from RNAflux results (**g**, **k**) not included in the respective pyTrance cluster.

**Extended Data Fig. 7:**
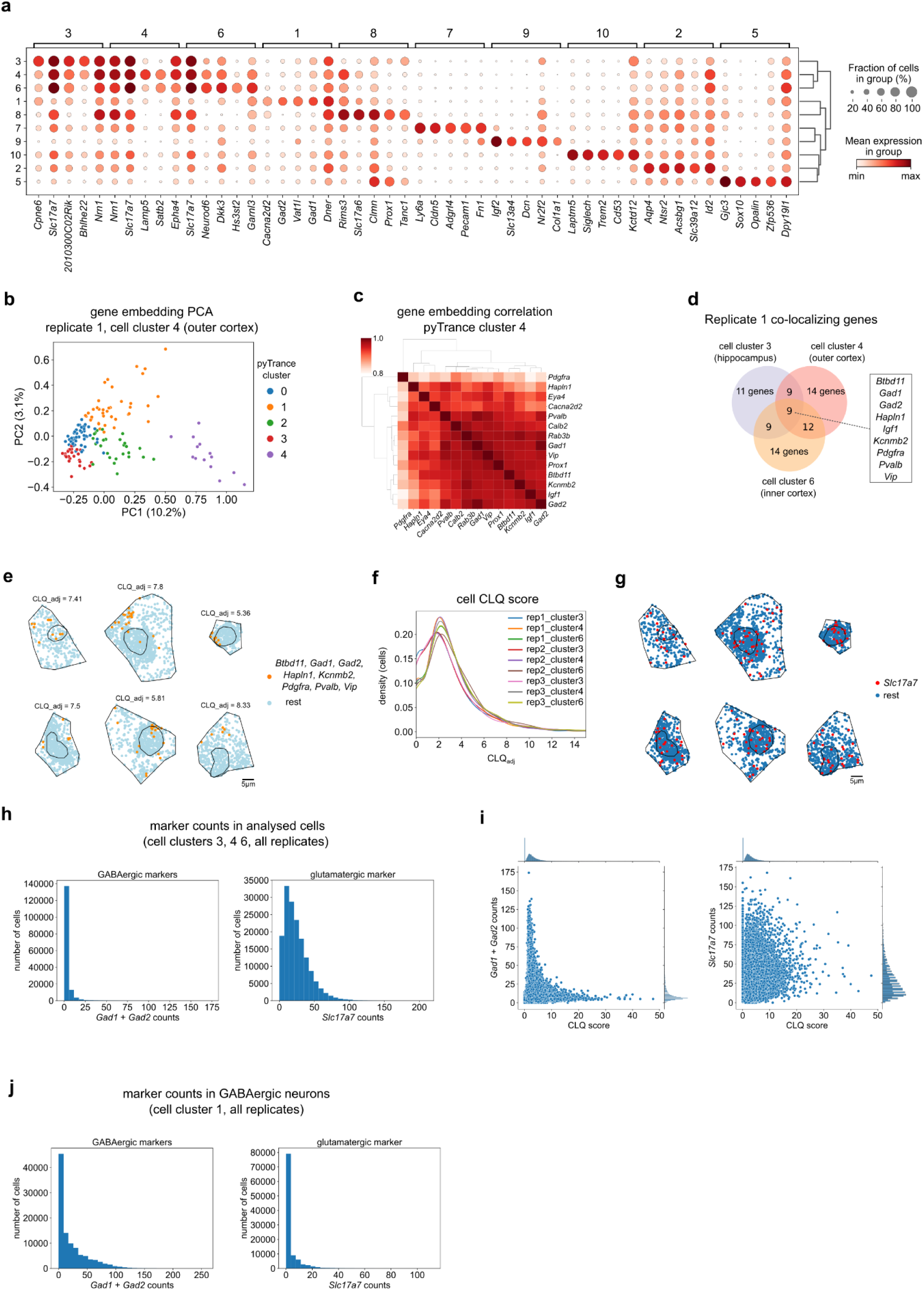
Extended mouse brain analysis results. **a**, Differentially expressed genes across the clusters of replicate 1 provided with the data. **b**, PCA of the gene embeddings computed with pyTrance on cell cluster 4 of replicate 1. **c**, Pairwise Pearson correlation matrices of gene embeddings of pyTrance cluster 4. **d**, Agreement between pyTrance clusters computed on different cell clusters of the same replicate. **e**, Single-transcript distributions highlighting the genes predicted to co-localize. Cells were sampled randomly from the top-scoring 10%. **f**, Kernel density estimations of the CLQ score distributions of *Gad1, Gad2, Hapln1, Kcnmb2* over the different cell clusters analysed. **g**, Single-transcript distributions highlighting *Slc17a7* in the same cells as in **e**. **h**, Count distributions of GABAergic (left) and glutamatergic neuron markers (right) in the cells analyzed with pyTrance. **i**, CLQ score versus counts of two GABAergic neuron markers (*Gad1, Gad2*; left) and *Slc17a7* (right). **j**, Count distributions of GABAergic (left) and glutamatergic neuron markers (right) in the cell clusters with GABAergic markers upregulated, which were not analyzed with pyTrance.

**Extended Data Fig. 8:**
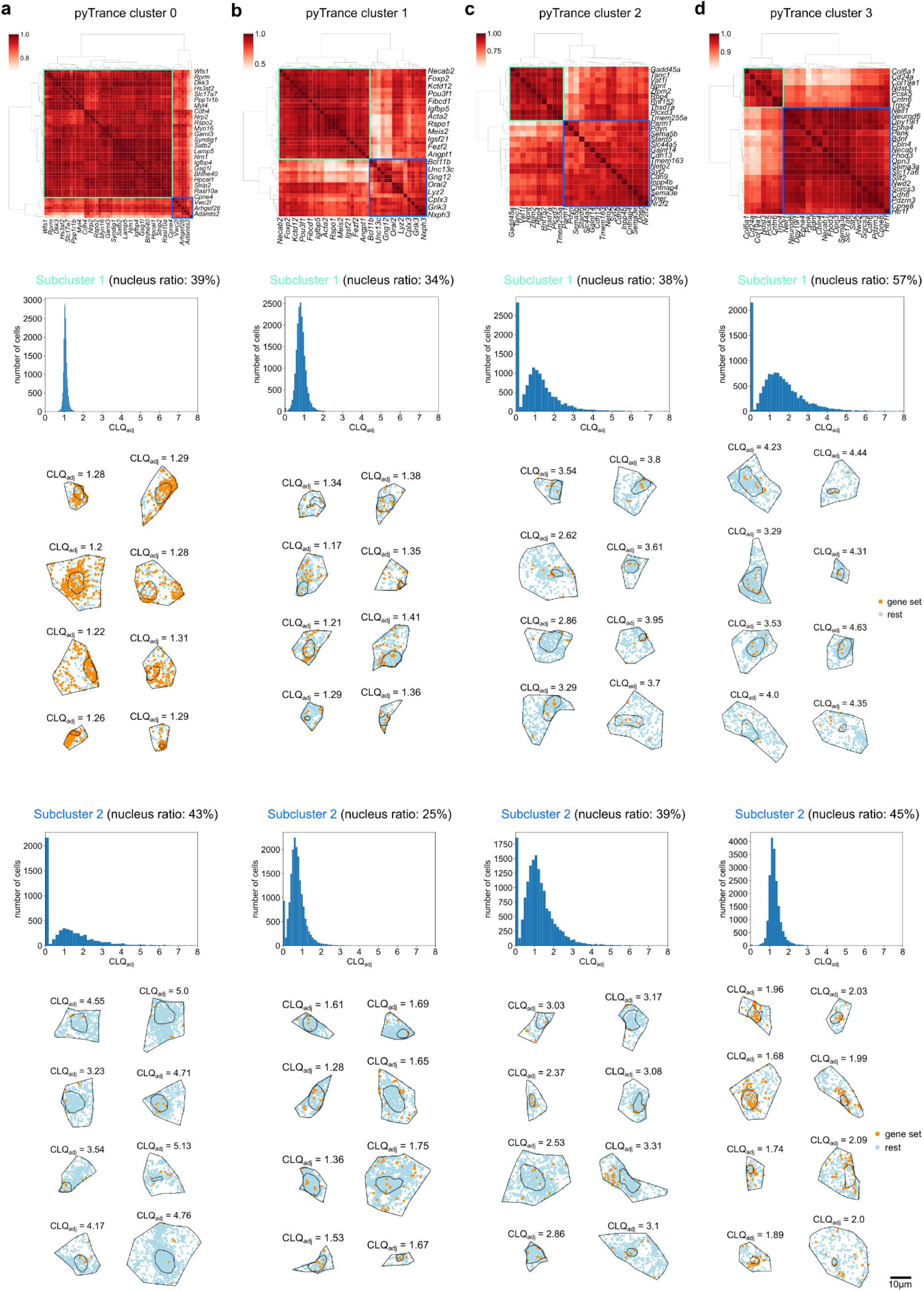
Predicted mouse brain co-localization subclusters (replicate 1, cell cluster 4). **a**-**d**, gene embedding correlation heatmap per pyTrance cluster with subclusters highlighted. **e**-**h**, CLQ score distributions over all cells for subcluster 1. **i**-**l**, Single-transcript distributions highlighting the respective genes per subcluster. **m**-**p**, CLQ score distributions over all cells for subcluster 2. **q**-**t,** CLQ score distributions over all cells for subcluster 2. Cells in **e**-**h** and **q**-**t** were sampled randomly from the top-scoring 10% per gene subcluster.

**Extended Data Fig. 9:**
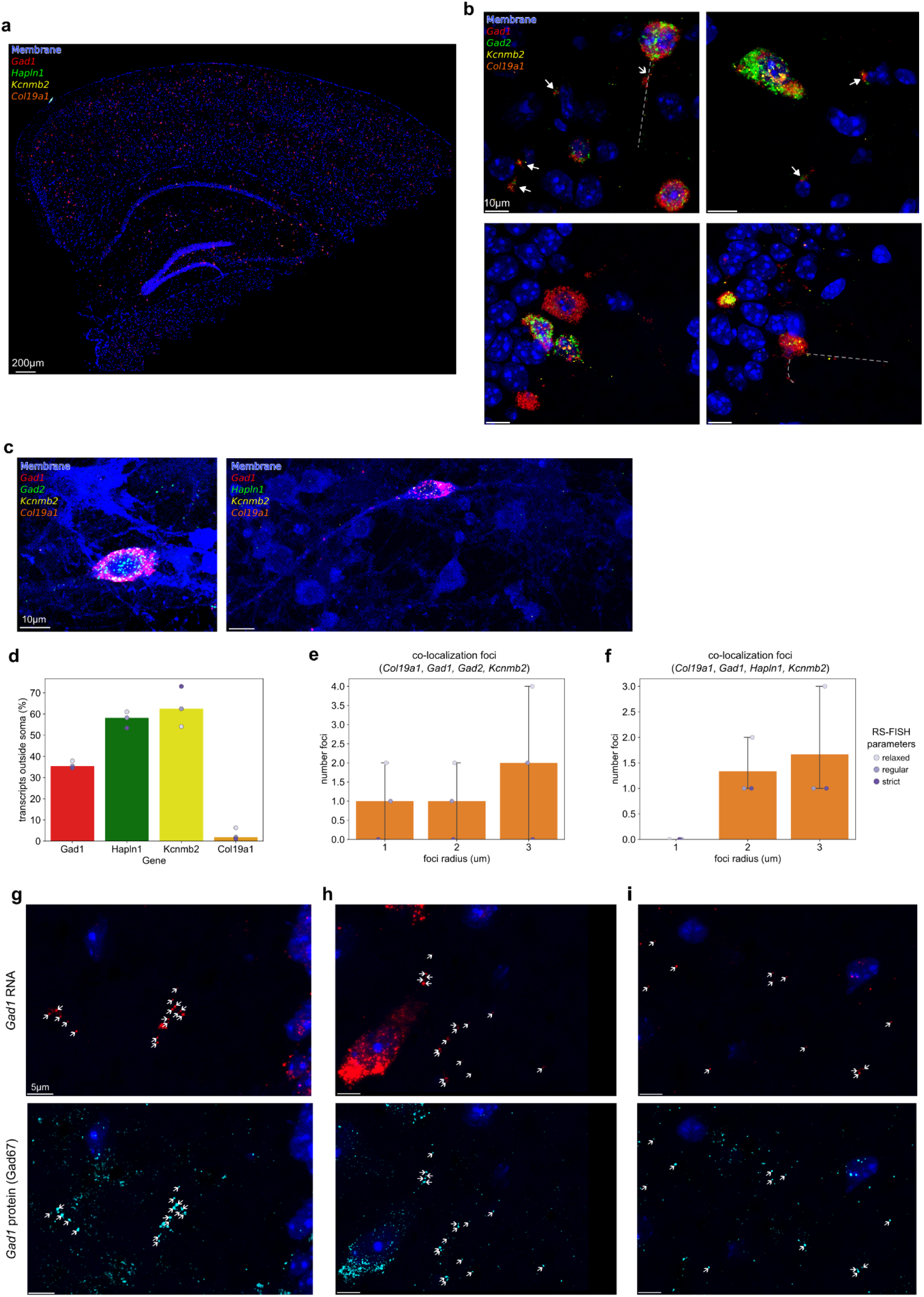
Validation of predicted RNA co-localization and *Gad1* RNA-protein co-localization. **a**, Full smFISH image of the second mouse brain section. **b**, Selected regions of interest on the first sections smFISH image. Closed-head arrows indicate co-localization of the three included candidate RNAs next to nuclei that show otherwise no expression, open-head arrows indicates foci where the RNAs co-localize on a projection, dashed lines indicate neuronal projections based on the RNA molecule arrangement. **c**, Example images of cultured neurons showing localization of RNAs to neuronal projections. **d**, Quantification of RNA localization based on three different parameter settings for molecule detection in brain section 2. **e**, **f**, **g,** Examples of *Gad1* RNA (top) and protein (bottom) co-localization in a third section. Arrows indicate positions of single RNA molecules that co-localize with proteins.

## Notes

### Competing Interest Statement

The authors have declared no competing interest.

